# Corticosterone-linked microglial activity underpins sexually dimorphic neuroplasticity after ketamine anesthesia

**DOI:** 10.1101/2025.10.19.683299

**Authors:** Alessandro Venturino, MohammadAmin Alamalhoda, Thomas Negrello, Kelly Jin, Cindy T.J. van Velthoven, Ryan John A. Cubero, Jake Yeung, Peter Koppensteiner, Bosiljka Tasic, Sandra Siegert

**Affiliations:** Institute of Science and Technology Austria (ISTA), Am Campus 1, 3400 Klosterneuburg, Austria; Allen Institute for Brain Science, Seattle, WA, USA; Laboratory of Neuroepigenetics, Brain Research Institute, Medical Faculty of the University of Zürich and Institute for Neuroscience, Department of Health Sciences and Technology, ETH Zürich, Zürich, Switzerland; Department of Computational Biology and Translation, Genentech Inc., 1 DNA Way, South San Francisco, CA 94080 USA

**Keywords:** microglia, sex, adulthood, ketamine anesthesia, corticosterone, plasticity, Fkbp5, visual cortex

## Abstract

Anesthesia recovery is critical for resuming normal physiological and neuronal functions; however, the mechanisms involved remain elusive. Here, we identify a female-selective corticosterone-mediated microglia-neuron interaction *in vivo* during ketamine anesthesia recovery, absent in males. This microglia-neuron interaction induces plastic and functional neuronal changes, as evidenced by increased spine density and mEPSC frequency, which is occluded upon microglia depletion. We show that this process is driven through upregulation of the stress-responsive co-chaperone *Fkbp5* mRNA and its protein, FKBP51, in female microglia. *Fkbp5*/FKBP51 is a key intermediary in a corticosteroid-induced stress response, and its involvement points towards a critical interface between endocrine signaling and microglia. Thus, to counteract the observed KXA-mediated corticosterone increase in the blood, we remove the primary source of corticosterone through adrenalectomy. Close microglia-neuron interaction was absent, but was reinstated after corticosterone injection. Our findings offer a new mechanism of microglia-mediated neuronal plasticity during anesthesia recovery, which is mediated through corticosterone, enhancing our understanding of sex differences in brain function.

## Introduction

Recovery from anesthesia is a fundamental yet complex process critical for resuming physiological functions. Ketamine distinguishes itself from other anesthetics due to its unique pharmacological properties as an NMDA receptor antagonist, preferentially targeting GABAergic inhibitory interneurons (*1*, *2*), and has been generally considered to comply to the principles of general anesthesia, which requires a drug-inducible, fully-reversible change of consciousness and cognition (*3*, *4*). However, repeated exposure to ketamine anesthesia reinstates juvenile-like plasticity in the primary visual cortex,mediated through alterations in the extracellular perineuronal nets (*5*). Moreover, ketamine anesthesia induces mild anxiety behavior phenotypes, interestingly, only in females (*6*), suggesting inherent sex differences in anesthesia recovery with neuronal consequences that extend beyond the immediate sex-dependent metabolic processing described for low-dose ketamine (*7*).

Ketamine, across different dosages, affects microglia (*5*, *8*–*10*), which are embedded within the neuronal network (*11*, *12*). Locally, microglia influence the synaptic machinery and neuronal firing properties by responding to environmental changes (*9*, *13*–*16*). Sex differences have been reported to influence microglia function (*8*, *17*–*20*), yet as immune-associated cells, microglia are underexplored in the emerging topics of sex differences across immune response (*21*, *22*) and neuronal activity (*23*, *24*).

Here, we investigate whether sex-specific microglia-neuron dynamics exist in a murine model subjected to ketamine-induced anesthesia. Only female mice exhibited a pronounced interaction between microglia and neurons during recovery, followed by increased synaptic density and enhanced neuronal firing properties. Mechanistically, we discovered that female microglia selectively upregulated the co-chaperone *Fkbp5*/FKBP51, which is a key intermediary in a corticosteroid-induced stress response (*25*–*27*). The selective hypothalamic activation and elevated corticosterone levels in blood plasma during the recovery phase in females shape the microglia-neuron interaction, highlighting a link between the endocrine and the brain-immune axis.

## Results

### Microglia-enabled neuronal network adaptation upon KXA recovery only in females

Anesthesia is typically divided into the initiation, deep anesthesia maintenance, and recovery stage (**Figure 1A**). We administered a single dose of ketamine-xylazine-acepromazine (KXA) anesthesia and performed immunostaining of the primary visual cortex (VISp) 4 hours later. We used Iba1 to identify microglia (*28*), the endosomal-lysosomal marker CD68 as a proxy for microglial reactivity (*29*), and vGluT2 staining as a reference to separate the cortical layers (**Figure S1A-B**). Only female microglia in cortical layers II/III, IV, and V/VI upregulated CD68 (**Figure S1A-C**). Similarly, the microglia 3D-morphology indicated a shift of the KXA female population to the saline condition in the latent space (**Figure S1D**), analyzed with an adapted morphOMICs (*8*) without bootstrapping. In contrast, KXA male microglia morphologies remained intermingled with saline (**Figure S1E**). The KXA female microglia density distribution shifted significantly in II-VI (**Figure S2**, **Methods**). In addition, CD68 relocated from the soma to the KXA female microglia process tips (**Figure S3A-B**) and also positively labeled for the perineuronal nets-staining *Wisteria floribunda agglutinin* (WFA, **Figure S3C-F**) (*5*). WFA selectively binds to the N-acetyl-galactosamine group of the chondroitin sulfate proteoglycans, which is part of the perineuronal nets amongst others in the cortical layer IV (*30*).

**Figure 1.**
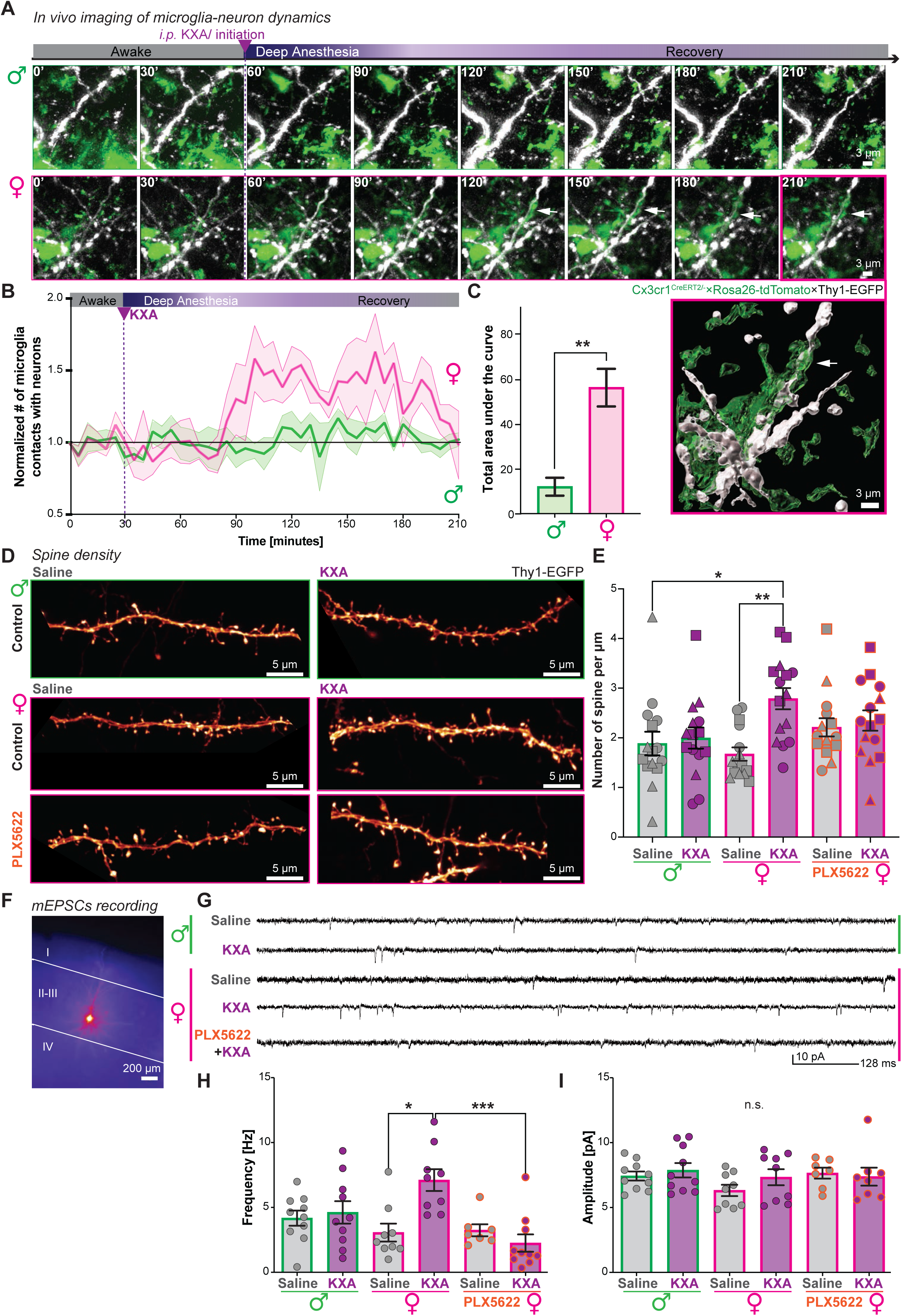
Female microglia interact with neurons, promoting spinogenesis and plasticity upon ketamine recovery. (**A-C**) Long-term *in vivo* 2-photon imaging in the primary visual cortex (VISp) through a cranial window of Cx3cr1^GFP/-^×Thy1-EGFP mice of both sexes spanning the phases of awake, deep anesthesia, and recovery after KXA (ketamine-xylazine-acepromacine) administration. (**A**) Sequential snapshots of microglia (green) and excitatory projection neurons (white). Top, males. Bottom, females. White arrow, prolonged microglia-dendrite contact. Magenta frame: 3D surface rendering of the 210-minute frame in females. Scale bar: 3 µm. (**B**) Normalized number of microglia and Thy1-EGFP neuronal process contacts over time in males (green) and females (magenta) represented as mean ± SEM confidence band. Dashed green line: KXA injection. 5 animals per condition. (**C**) Bar chart of the mean total area under the curves in (**B**). Mean ± SEM of 5 animals per condition. Unpaired t-test with Welch’s correction, **p < 0.01. (**D-E**) Spine density quantification in VISp, layer II/III in Thy1-EGFP mice 4 hours after saline or KXA injection. (**D**) Representative super-resolution images of dendritic processes. Top: males, middle: females, bottom: PLX5622-treated females for 1.5 weeks to deplete microglia. Scale bar: 5 µm. (**E**) Bar chart of the mean number of spines per µm with ± SEM. Each point, number of spines. 5 dendrites/animal. Symbols, different animals. 3 animals per condition. Kruskal-Wallis with selected Dunn’s multiple comparisons post hoc test, *p < 0.05, **p < 0.01. (**F-I**) mEPSC (miniature excitatory postsynaptic currents) recording from layer II/III VISp pyramidal neurons, 4 hours after saline or KXA injection in either males, females, or microglia-depleted females (PLX5622). (**F**) Representative epifluorescence image of a biocytin-filled pyramidal neuron after mEPSC recording. Scale bar: 200 µm. (**G**) Representative recorded mEPSC example traces. (**H-I**) Bar charts of mean mEPSC frequency (**H**) and amplitude (**I**) with SEM. Each dot, a recorded neuron. 2-4 cells/animal. 3 animals/condition. Kruskal-Wallis with selected Dunn’s multiple comparisons post hoc test, *p < 0.05, ***p < 0.001, ^ns^p > 0.05, not significant.

To verify the sex-specific effect and to identify at which anesthesia stage female microglia start to respond, we imaged microglial dynamics *in vivo* in the VISp. We used the Cx3cr1^CreERT2/het^×Ai9-tdTomato mouse model, crossed with Thy1-EGFP (*31*–*34*), to visualize sparsely labeled pyramidal neuronal processes reaching cortical layers I and II. During the baseline recording for 30 minutes, microglial processes infrequently contacted the neuronal processes and spines, with no differences noted between the sexes (**Figure 1A**). After the *i.p.* injection of KXA, microglial contacts remained similarly active as during baseline for approximately 60 minutes during maintenance (**Figure 1B**). Remarkably, one hour following the single dosage of KXA injection, only female microglia established significantly more contacts with the labeled pyramidal neuronal processes and spines (**Figure 1C**, **Supplementary Videos 1-2**). This interaction gradually diminished after three hours of *in vivo* recording (**Figure 1B**), as the animal slowly awoke.

To determine whether the close interaction of microglial processes with pyramidal cells correlates with changes in dendritic processes (*16*, *35*, *36*), we quantified spine density in the second-order dendrites four hours after KXA using super-resolution imaging (**Figure 1D**). Only females subjected to KXA showed an increase in spine density compared to saline conditions, while the number of spines in males remained unchanged (**Figure 1E**). These results also aligned with the recorded spontaneous miniature excitatory synaptic currents (mEPSCs) from layer II/III pyramidal cells in slice recordings (**Figure 1F-G**). The mEPSC frequency increased significantly only in females (**Figure 1H**), while the amplitude remained unchanged across both sexes (**Figure 1I**). To establish the relevance of microglia in this process, we provided mouse chow containing CSF1R blocker PLX5622 to mice 1.5 weeks before the KXA experiment. This treatment achieved approximately 80% microglia depletion (**Figure S4**), and it abolished the effects of increased spine number and mEPSC frequency under the same experimental design (**Figure 1D-I**), supporting the notion of female-selective microglia-enabled neuroplasticity.

### KXA affects genes associated with the glucocorticoid pathway

To obtain mechanistic insights into the pathway driving the microglia-mediated response, we performed single-nucleus multiome sequencing of the entire cell population in female VISp two hours after KXA induction (**Figure 2A**). The total dataset consisted of 36,701 cells after quality control and removal of low-quality cells (**Figure S5A-D**). After batch-corrected normalization, we assigned the clusters to individual cell types (**Figure 2B**, **Figure S5E**) and verified that each sample had a comparable cell-type composition (**Figure S5F-G**). Each cell type exhibited a similar ratio of saline and KXA-containing cells (**Figure S5H**). The overall effect of KXA across the entire population did not produce a prominent cluster or shift among the cell populations. Furthermore, the cell type distribution across the four samples and within the condition remained similar, indicating no loss or proliferation of cells, particularly for microglia and astrocytes.

**Figure 2.**
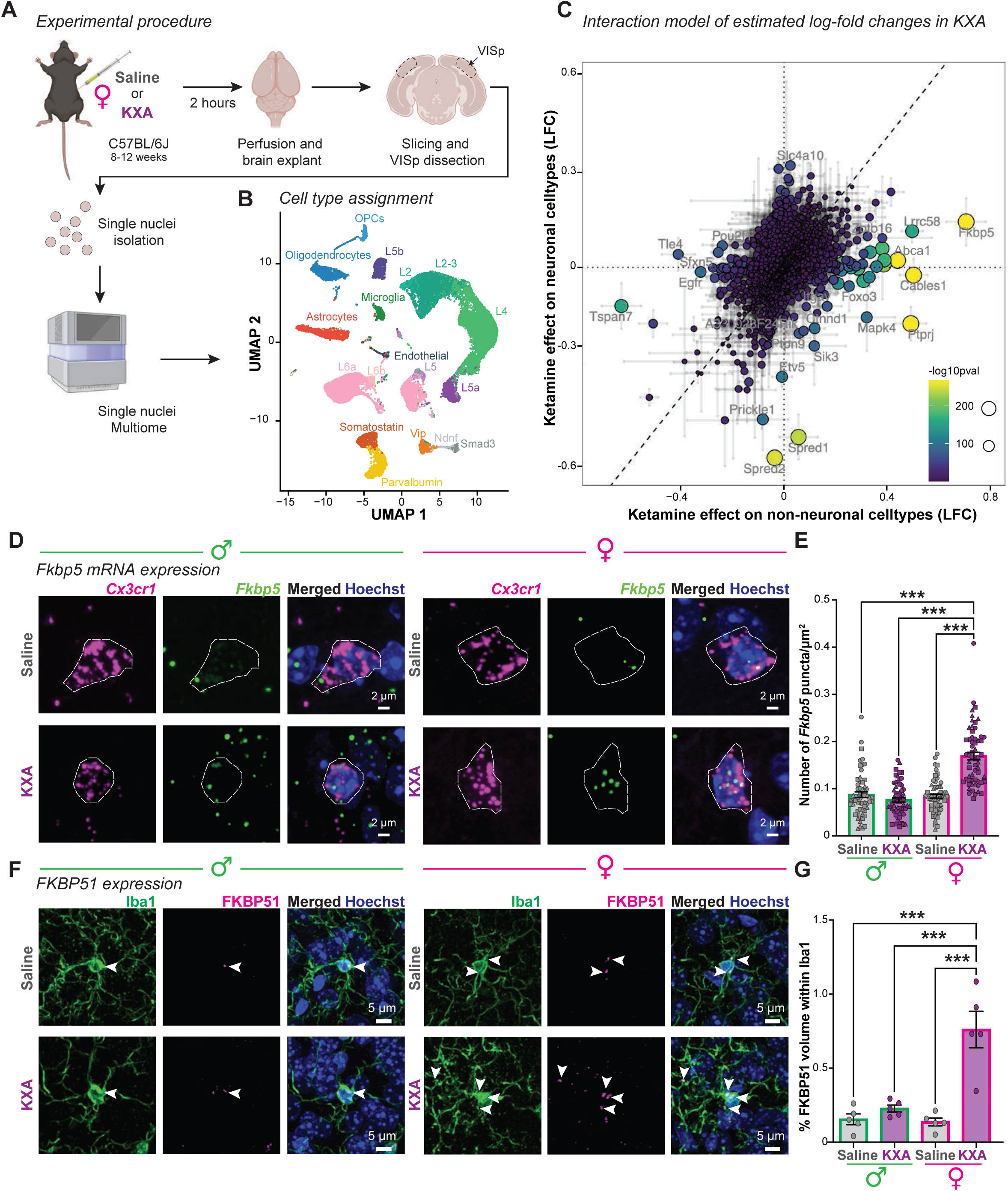
Microglia upregulate the stress-mediator *FKBP5*/FKBP51. (**A**) Experimental strategy for single-nuclei multiome sequencing. Primary visual cortex (VISp) of female mice microdissected 2 hours after saline or KXA (ketamine-xylazine-acepromazine) treatment. (**B**) UMAP plot of cell type assignment. (**C**) Log fold change (LFC) estimates from KXA treatment on non-neuronal (astrocytes and microglia) *versus* neuronal cell types. Size and yellow intensity of points are proportional to the -log_10_(p-value) from ANOVA for interaction between ketamine effect and astrocytes/microglia cell type (see Methods). Error bars: 90% confidence intervals inferred from KXA effect on non-neuronal or neuronal cell types. Genes along the dashed line have equal LFC and zero interaction effect. (**D-E**) Fluorescence *in situ* hybridization for male and female mice 2 hours after saline or KXA injection in VISp. (**D**) Example high-magnification images of mRNA probes against *Fkbp5* (green) and *Cx3cr1* (magenta, microglia), counterstained with the nuclei-dye Hoechst (blue), which provides the nucleus contour (white dashed line). Scale bar: 2 µm. (**E**) Bar chart of the mean *Fkbp5* mRNA puncta within the Hoechst contour with SEM. Each dot represents one microglia contour, 20 cells per animal, 3 animals/condition. Kruskal-Wallis with selected Dunn’s multiple comparisons post hoc test, ***p < 0.001. (**F-G**) FKBP51 protein expression 4 hours after saline or KXA injection in the VISp, layers III to V of males and females. (**F**) Example immunofluorescence images for Iba1 (green) and FKBP51 (magenta), counterstained with the nuclei-dye Hoechst (blue). White arrowheads: FKBP51 localization within Iba1^+^-microglia. Scale bar: 5 µm. (**G**) Bar chart of the mean percentage of FKBP51 volume within microglia with SEM. Each dot, one animal, 5 animals/condition. Two-way ANOVA with selected Tukey’s multiple comparisons post hoc test, ***p < 0.001.

Next, we identified the 2000 most differentially regulated genes in the global transcriptome (**Figure 2B**). We developed an interaction model to estimate log-fold changes from KXA treatment on non-neuronal astrocyte-microglia population *versus* neuronal cell types. Most genes had a small effect size and remained unaffected by KXA (**Figure 2C**). Specifically in neurons, only *Spred1* and *Spred2* were downregulated, which are associated with the MAP kinase cascade (*37*, *38*). In contrast, non-neuronal cells significantly upregulated *Abca1*, *Cables1*, *Ptprj*, and *Fkbp5*. A common theme among these genes is their relationship to glucocorticoid either in its production through cholesterol (*Abca1*) (*39*, *40*) and lipid metabolism (*Ptprj*) (*41*) or in its regulation (*Cables1*, *Fkbp5*) (*42*, *43*). Specifically, *Fkbp5* and its protein, the FK506-binding protein 51 (FKBP51), are essential stress response regulators (*25*–*27*), which tightly co-regulate the glucocorticoid receptor expression and the sensitivity of a cell to detect the stress hormone corticosterone (*26*, *27*, *43*, *44*).

To verify the selective upregulation of *Fkbp5* mRNA in female KXA microglia, we performed fluorescence *in situ* hybridization and quantified the number of *Fkbp5* puncta within the Hoechst nuclei contour using cell-type-specific probes in KXA-treated and non-treated saline neural tissue from both sexes. Only female *Cx3cr1^+^-*microglia exposed to KXA showed an increase in *Fkbp5* puncta. This effect did not occur in males (**Figure 2D-E**). Furthermore, this upregulation was specific to microglia, as S100β^+^-astrocytes and NeuN^+^-neurons displayed no significant differences between saline and KXA conditions (**Figure S6**). Notably, NeuN^+^-neurons from females exhibited a significantly lower abundance of *Fkbp5* at baseline than males, which remained unaffected upon KXA. Finally, we evaluated whether the upregulation of *Fkbp5* also translated to upregulation of the FKBP51 protein. Like *Fkbp5*, only female microglia upregulated FKBP51 four hours after KXA exposure (**Figure 2F-G**), suggesting that *Fkbp5*/FKBP51 contribute to the female microglial response during KXA recovery.

### Fkbp5 links to the female microglia-enabled effect on neuronal plasticity

*Fkbp5*/FKBP51 is an interesting drug target for pharmaceutical strategies due to its predicted gene-environment interactions at the human *FKBP5* locus in various mental disorders (*27*, *45*). Selective antagonists of the FK506-binding protein 51 by induced fit (including SAFit2) have been developed with a high binding affinity for FKBP51 (*45*, *46*). We injected SAFit2 with either KXA or saline and analyzed the overall effect on CD68 expression in microglia four hours later (**Figure 3A**). Inhibiting FKBP51 prevented KXA-mediated CD68 upregulation (**Figure 3B-C**) and PNN accumulation within the microglial CD68 compartments (**Figure S7A-C**). SAFit2 injected with saline did not directly affect microglia morphology and intermingled with control saline phenotypes. In contrast, SAFit2 and KXA partially mitigated the morphological shift observed for the KXA female microglia across the cortical layers (**Figure S8A-C**).

**Figure 3.**
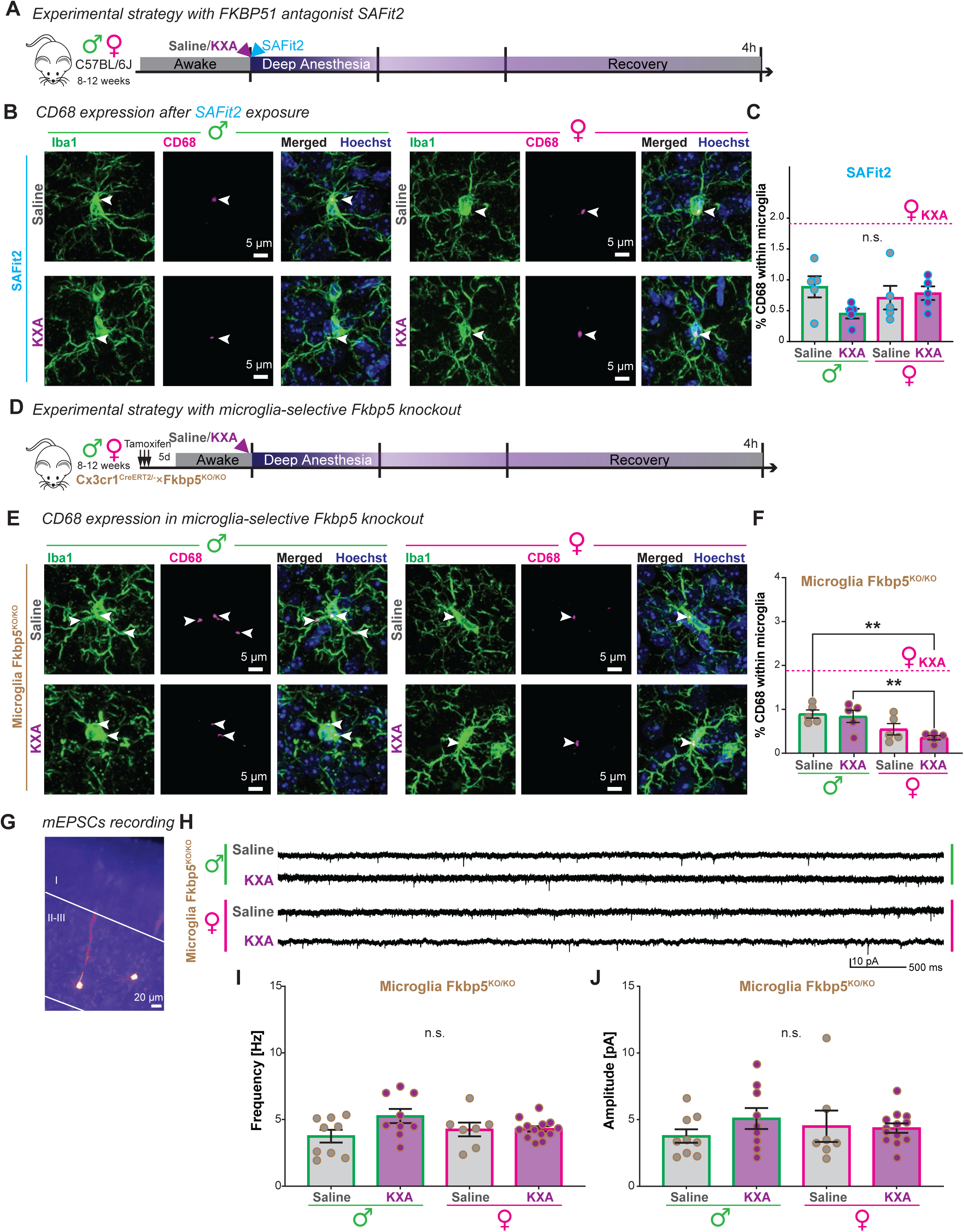
SAFit2 treatment or microglia-specific *Fkbp5* knockout prevents female microglia response and neuronal plasticity. (**A-C**) Experimental strategy for antagonising FKBP51 with SAFit2 after injection of saline or KXA (ketamine-xylazine-acepromazine). (**B-C**) CD68 quantification in primary visual cortex (VISp), layer III-V of males and females, 4 hours after saline or KXA with SAFit2. (**B**) Representative immunostainings for Iba1 (green) and CD68 (magenta), counterstained with the nuclei-dye Hoechst (blue). Arrow, CD68 localisation within microglia. Scale bar: 5 µm. (c) Bar chart of the mean percentage of CD68 volume within microglia with SEM. Each dot, one animal. 5 animals/condition. Dashed line, reference value of female KXA without SAFit2 from **Figure S1C**. Two-way ANOVA. ^ns^p > 0.05, not significant. (**D-J**) Consequences of microglia-selective *Fkbp5*-knockout experiment using a tamoxifen-inducible Cx3cr1^CreERT2/-^ ×Fkbp5^KO/KO^ reporter mouse line (Microglia Fkbp5^KO/KO^). (**D**) Experimental strategy. 3 consecutive tamoxifen injections, starting 8 days before performing the procedure. (**E-F**) CD68 quantification in primary visual cortex (VISp), layer III-V of males and females, 4 hours after saline or KXA. (**E**) Representative images of immunostainings for Iba1 (green) and CD68 (magenta), counterstained with the nuclei-dye Hoechst (blue). Arrow, CD68 localisation within microglia. Scale bar: 5 µm. (**F**) Bar chart of the mean percentage of CD68 volume within microglia with SEM. Each dot, one animal. 5 animals/condition. Dashed line, reference value of female KXA in C57BL/6J from **Figure S1C**. Two-way ANOVA with selected Tukey’s multiple comparisons post hoc test, **p < 0.01. (**G-J**) mEPSC (miniature excitatory postsynaptic currents) recording from layer II/III VISp pyramidal neurons, 4 hours after saline or KXA injection in males or females. (**G**) Representative epifluorescence image of a biocytin-filled pyramidal neuron after mEPSC recording. Scale bar: 20 µm. (**H**) Representative recorded mEPSC example traces. (**I-J**) Bar charts of mean mEPSC frequency (**I**) and amplitude (**J**) with SEM. Each dot, a recorded neuron. 2-4 cells/animal. 3 animals/condition. Two-way ANOVA, ^ns^p > 0.05.

Since SAFit2 affects all cells that express FKBP51, including neurons and astrocytes (**Figure S6**), we chose to examine a microglia-selective Fkbp5 knockout mouse line (*47*), Cx3cr1^CreERT2/het^×Fkbp5^flox/flox^, for KXA effects (**Figure 3D**). After confirming the selective FKBP51 loss in microglia following the tamoxifen-induced knockout (**Figure S9A-B**), we verified the lack of KXA-mediated CD68 level increase (**Figure 3E-F**) and the containment of WFA staining (**Figure S9F-G**), suggesting that the selective knockout of *Fkbp5* hindered the targeted microglial response upon KXA recovery. Notably, the microglial density remained unaffected, excluding macrophage infiltration (**Figure S9D-E**). Finally, to test effects on neuronal plasticity, we measured the frequency and amplitude of mEPSC in layer II/III pyramidal cells after saline or KXA exposure in both males and females (**Figure 3G-H**). In neither condition did we observe changes in frequency (**Figure 3I**) or amplitude (**Figure 3J**).

### Corticosterone as a driver of sex-specific microglia-mediated effects

*Fkbp5*/FKBP51 upregulation in female microglia during KXA recovery and the lack of a microglial response upon *Fkbp5* knockout and FKBP51 inhibition suggest a link between the body periphery and neuronal network adaptation mediated via the endocrine steroid hormone corticosterone, the only glucocorticoid in mice (*48*). The hypothalamic paraventricular nucleus (PVN) is one of the key regions that regulates corticosterone levels by releasing corticotropin-releasing hormone (CRH). The anterior pituitary gland converts CRH into adrenocorticotropic hormone (ACTH), which stimulates the adrenal glands to release corticosterone (*39*, *49*).

To systematically establish the connection between corticosterone and the microglia-mediated effects, we used the Targeted Recombination in Active Population (TRAP-cFos) mouse line (*50*), crossed with the Ai9 reporter line. We trapped the neurons with 4-OH-tamoxifen starting 30 minutes after KXA to capture c-Fos-dependent neuronal activity during the recovery procedure between both sexes (**Figure 4A-B**). Remarkably, we found a significantly higher number of trapped neurons in the female PVN than in males (**Figure 4B**). To verify whether this translates into increased corticosterone levels, we collected blood plasma after 30 or 120 minutes from saline or KXA-injected mice (**Figure 4A**). After 30 minutes, corticosterone levels remained comparable between saline and KXA injections, with no differences between the sexes (**Figure 4C**). Remarkably, only in females did we find a three-fold increase of corticosterone 120 min after KXA, suggesting an intrinsic, sex-dependent response during the KXA recovery.

**Figure 4.**
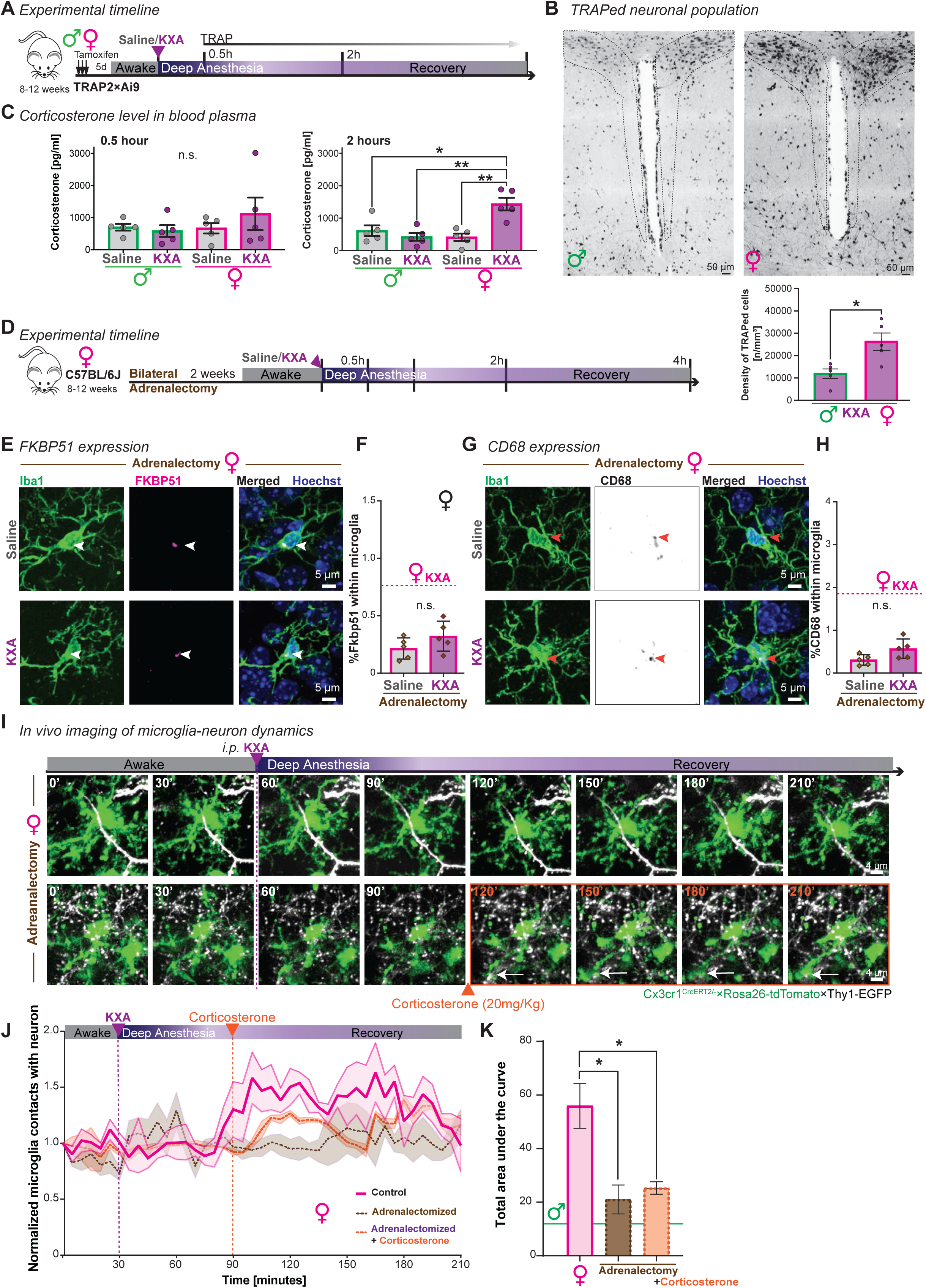
Ketamine-induced corticosterone drives female microglia-neuron interaction. (**A**) Experimental timeline after saline or KXA (ketamine-xylazine-acepromacine) injection for TRAP-based neuronal activity tagging in TRAP2×Ai9 mouse line and systemic blood collection for corticosterone quantification in C57BL/6J. (**B**) Representative images of TRAPed neurons 30 min after KXA injection in the paraventricular nucleus (PVN, dashed line, coronal brain slice) in males (left) and females (right) in TRAP2×Ai9 mice. Scale bar: 50 µm. Below, bar chart of the mean density of TRAPed neurons in the PVN with SEM. Each dot, one animal. 5 mice/condition. Unpaired t-test with Welch’s correction, *p < 0.05. (**C**) Corticosterone level quantification in blood plasma isolated from males (green) and females (magenta) 30 minutes (left) or two hours (right) after saline or KXA treatment. Each dot, one animal. 5 mice/condition. Two-way ANOVA with selected Tukey’s multiple comparisons post hoc test shown, *p < 0.05, **p < 0.01. (**D**) Experimental timeline for bilateral adrenalectomy in females. After 2 weeks of recovery, injection of saline or KXA. (**E-H**) Immunostainings and quantification in the primary visual cortex (VISp), layer III-V of males and females, 4 hours after saline or KXA treatment. (**E**, **G**) Representative images of immunostainings for Iba1 (green) and FKBP51 (magenta) (**E**) or CD68 (grey) (**G**), counterstained with the nuclei-dye Hoechst (blue). Arrow, localisation within microglia. Scale bar: 5 µm. (**F**, **H**) Bar chart of the mean percentage of FKBP51 (**F**) and CD68 (**H**) volume within microglia with SEM. Each dot, one animal, 5 animals/condition. Dashed line, reference value of female KXA for FKBP51 (Figure 2G) and CD68 (**Figure S1C**). Welch’s t-test. ^ns^*p* > 0.05, not significant. (**I-K**) Long-term *in vivo* 2-photon imaging in the primary visual cortex (VISp) through a cranial window of adrenalectomized C57BL/6J females spanning the phases of awake, deep anesthesia, and recovery after KXA administration. (I) Sequential snapshots of microglia (green) and excitatory projection neurons (white). Top, only adrenalectomized females. Bottom, with *i.p.* corticosterone injection at 120 minutes (orange). White arrow, prolonged microglia-dendrite contact. Scale bar: 4 µm. (J) Normalized number of microglia and Thy1-EGFP neuronal process contacts over time in females (magenta, see Figure 1B), adrenalectomized females (purple, n=3), and adrenalectomized females injected with corticosterone 60 minutes after KXA injection (orange, n=3) represented as mean ± SEM confidence band. Dashed lines: KXA (green) and corticosterone (orange) injections. (**K**) Bar chart of the mean total area under the curves with ± SEM from (**J)**. Green line, reference value of male from Figure 1C. Mean ± SEM of 3 - 5 animals per condition. Brown-Forsythe ANOVA test with Dunnett’s T3 multiple comparisons test, *p < 0.05.

The adrenal glands are the primary source of corticosterone (*49*). We performed bilateral adrenalectomy to prevent the KXA-mediated rise in corticosterone (**Figure 4D**). After the surgery, the animals were exposed to low corticosterone dosage in the drinking water, avoiding alterations in the circadian rhythm (*51*). To confirm that we prevented the FKBP51 increase in microglia, we analyzed the expression level within microglia in adrenalectomized females. Following KXA exposure, those females exhibited significantly less FKBP51 expression within microglia than the non-adrenalectomized animals (**Figure 4E-F**, **Figure S10A**). Furthermore, microglia expressed lower levels of CD68 (**Figure 4G-H**, **Figure S10B**). Lastly, to link microglia-mediated FKBP51-corticosterone to neuronal interaction, we performed cranial window surgery in Cx3cr1^CreERT2/het^×Ai9×Thy1-EGFP adrenalectomized females and imaged microglia-neuron interaction *in vivo*. In adrenalectomized females, microglia did not interact closely with the pyramidal neurons (**Figure 4I-K**), as previously observed (**Figure 1A-C**). Given the known increase of *Fkbp5* and corticosterone after two hours, we injected 20 mg/Kg of corticosterone 90 minutes after KXA injection in adrenalectomized females to stimulate an extra corticosterone surge (*52*, *53*), such as observed under KXA (**Figure 4C**, **Supplementary Video 3**). This corticosterone injection reinstated the *in vivo* microglia-neuron interaction (**Figure 4I-K**, **Figure S10C**, **Supplementary Video 4**), establishing a direct link between corticosterone-associated blood levels and microglia interaction with neurons via *Fkbp5*/FKBP51.

## Discussion

Our study uncovers significant sex-specific differences in neuronal adaptation during ketamine anesthesia recovery, enabled by microglia (**Figure S11**). Specifically, female mice showed enhanced microglial activity in response to corticosterone elevation following ketamine anesthesia, facilitated through the *Fkbp5*/FKBP51 pathway. This response increased synaptic density and neuronal plasticity, underscoring a critical role for microglia in female neural adaptation— a process not observed in males at the selected time point.

These findings contribute to a growing body of evidence that recognizes sex-specific differences in brain and immune responses, the latter describing differences in susceptibility to infection and autoimmune diseases (*21*, *22*). Previous research has often overlooked these distinctions, treating them as minor (*24*); however, recent advancements highlight the substantial impact of sex on physiological and behavioral responses (*23*).

Our research positions the glucocorticoid pathway as a mediator of these sex-specific recovery effects. Through FKBP51, microglia modulate the effects of corticosterone and its impact on the neuronal network. This modulation emphasizes the complexity of glucocorticoid action in the context of neural recovery.

The discovery that corticosterone suffices to trigger these pathways implicates the stress response as an integral part of the anesthesia recovery dynamics. Unlike studies focusing on ketamine as an antidepressant therapy, our work emphasizes the physiological effects post-anesthesia and posits microglia as key intermediaries. In the future, investigations should focus on the interplay between ketamine and other cellular actors, like astrocytes, which have shown interconnected responses in neural plasticity (*54*).

In conclusion, our study highlights significant sex-specific microglia-neuron interplay during anesthesia recovery in the adult brain, mediated by the glucocorticoid pathway. This emphasizes that circulating sex hormones are not the sole factor accounting for differences (*24*) and that exploring differences expands our knowledge on how the brain adapts to pharmacological interventions. Our findings underline that microglia are a relevant interface between the endocrine stress response and the brain-immune cell system.

## Materials and methods

### Animals

If not otherwise indicated, we used adult mice (8-12 weeks) of both sexes. C57BL/6J (Cat#000664), Cx3Cr1^CreERT2^ (B6.129P2(C)-Cx3cr1tm2.1(cre/ERT2)Jung/J, Cat#020940, always used heterozygous) (*55*), Ai9 (B6;129S6-Gt(ROSA)26Sortm9(CAG-tdTomato)Hze/J, Cat#007909) (*56*), Thy1-GFP (Tg(Thy1-EGFP)MJrs/J, Cat#007788) (*57*), and TRAP2 ((Fostm2.1(icre/ERT2)Luo/J), Cat#030323, crossed with Ai9, (*50*)) were purchased from The Jackson Laboratories. FKBP5^flox/flox^ mice were kindly provided by Mathias V. Schmidt (*47*). All mice were housed in the ISTA Preclinical Facility with a 12-hour light-dark cycle. Food and water were provided *ad libitum*. The estrous phase was not systematically determined, except for the multiome analysis. All animal procedures are approved by the Bundesministerium für Wissenschaft, Forschung und Wirtschaft (bmwfw) Tierversuchsgesetz 2012 (TVG 2012), BGBI. I Nr. 114/2012, idF BGBI. I Nr. 31/2018 under the numbers 66.018/0005-WF/V/3b/2016, 66.018/0010-WF/V/3b/2017, 66.018/0025-WF/V/3b/2017, 66.018/0001_V/3b/2019, 2020-0.272.234.

### Drugs

If not otherwise indicated, mice received *intraperitoneal* (*i.p.*) injections on weekdays in the morning. Since circadian cues might affect the experimental readout, we performed the experiments in the morning when the peak levels of glucocorticoids are reduced (*58*, *59*).

#### Ketamine-xylazine-acepromazine (KXA)

We combined ketamine (100 mg/Kg, MSD Animal Health, Cat #A137A01) with xylazine (10 mg/Kg, Livisto, Cat#7630120), which prevents ketamine-induced muscle rigidity, and acepromazine (3 mg/Kg, VANA GmbH, Cat#18F211), a phenothiazine tranquilizer (KXA) (*60*), solubilized in physiological saline solution (0.9% (w/v) NaCl, Fresenius Kabi Austria, Cat#19MIA700). The solution was always freshly prepared to avoid pH fluctuations. As a control, we injected saline solution with the same volume as KXA.

#### Tamoxifen

Adult Cx3Cr1^CreERT2/*het*^×Ai9 and Cx3Cr1^CreERT2/*het*^×FKBP5^flox/flox^ mice were injected with 150 mg/Kg tamoxifen (Sigma, Cat#T5648, Lot WXBD2299V) dissolved in corn oil (Sigma, Cat#MKCH1635) for three consecutive days. Experiments started 5 days after the last injection.

#### 4-hydroxy-tamoxifen

Injections were performed as previously described (*61*). In brief, TRAP2-Ai9 mice were injected with 50 mg/Kg 4-hydroxytamoxifen (Sigma, Cat#H6278) in corn oil. The day before the experiment, 10 mg of 4-hydroxytamoxifen powder was solubilized in 1000 µL of pure ethanol. The solution was divided into 150 µL aliquots and stored overnight at -20°C. On the day of the experiment, the aliquots were suspended in 150 µL of corn oil, creating two phases with ethanol (top) and corn oil (bottom). The suspensions were centrifuged using a Speed Vac (ThermoFisher, Cat#SPD210) for 15 minutes at 40°C until the top phase was evaporated entirely. The bottom phase, containing corn oil and 4-hydroxytamoxifen, was injected into the animals.

#### PLX5622

The animals received *ad libitum* chow 1.5 weeks before KXA injection, containing 1200 mg/Kg PLX5622 (DC Chemicals, Cat#DC21518), incorporated into the food pellets (Sniff, Cat#S4865-E012). Mice were kept on this diet for the entire experiment (*62*, *63*).

#### Corticosterone

Corticosterone synthetic powder (≥92% purity, Sigma, Cat#C2505) was solubilized in 50 µL DMSO (Sigma, Cat# D8418) and subsequently diluted in a 1:5 ratio with saline to reduce toxicity (*64*). During the two-photon imaging session, adrenalectomized Cx3Cr1^CreERT2/het^×Ai9×Thy1-GFP mice were injected with 20 mg/Kg corticosterone (*65*).

#### SAFit2

SAFit2 (MedChem Express, Cat#HY-102080) was dissolved in 90% (v/v) corn oil and 10% (v/v) ethanol, and 20 mg/Kg (*66*) was injected.

### Anesthesia induction

After the animals received the KXA injection, their eyes were treated with eye ointment (Oleo Vital) to prevent corneal dehydration, and they were maintained at 37°C throughout the procedure. The achievement of deep anesthesia was confirmed based on the following parameters: 1. Absence of the toe pinch reflex 10 minutes after induction. 2. Decrease in respiratory frequency. 3. No responses to noxious stimuli. 4. Flaccid paralysis. 5. Absence of whisker movement (*67*).

### Immunostaining

#### Brain tissue preparation

If not otherwise indicated, the brain was dissected four hours after the drug treatment, exceeding the KXA half-life of 2-3 h (*68*). The animal was quickly anesthetized with isoflurane (Zoetis, Cat#6089373) and secured to the perfusion plate. The chest was opened to expose the heart. The left ventricle was cannulated, and the *inferior vena cava* was cut. The animals were initially perfused with 20 mL of phosphate-buffered saline (PBS) containing heparin (100 mg/L, Sigma, Cat#H0878), followed by 20 mL of 4% (w/v) paraformaldehyde (PFA, Sigma, Cat#P6148) in PBS using a peristaltic pump (Behr, Cat#PLP 380, speed: 25 rpm). The animal was decapitated, the brain explanted, and post-fixed in 4% (w/v) PFA/PBS overnight (16h) at 4°C. Then, the tissue was washed in PBS and stored in PBS with 0.025% (w/v) sodium azide (VWR, Cat#786-299) at 4°C. For cryoprotection, the tissue was transferred to 30% (w/v) sucrose (Sigma, Cat#84097) in PBS and incubated overnight at 4°C. The brains were kept at -70°C in 30% (w/v) sucrose for long-term storage. If not otherwise indicated, the brain was sliced into 100 μm coronal sections on a vibratome (Leica VT 1200S).

#### Immunohistochemistry

The brain slices were incubated in a blocking solution containing 1% (w/v) bovine serum albumin (Sigma, Cat#A9418), 5% (v/v) Triton X-100 (Sigma, Cat#T8787), 0.5% (w/v) sodium azide (VWR, Cat#786-299), and 10% (v/v) serum (either goat, Millipore, Cat#S26, or donkey, Millipore, Cat#S30) for 1 hour at room temperature on a shaker. Afterward, the samples were immunostained with primary antibodies diluted in an antibody solution containing 1% (w/v) bovine serum albumin, 5% (v/v) Triton X-100, 0.5% (v/v) sodium azide, and 3% (v/v) goat or donkey serum for 48 hours on a shaker at room temperature. The following primary antibodies were used: rat α-CD68 (BioRad, Cat#MCA1957, Lot 155083, 1:250); goat α-Iba1 (Abcam, Cat#ab5076, Lot1014660-1, 1:250); rabbit α-Iba1 (GeneTex, Cat#GTX100042, Lot 44200, 1:750); rabbit α-FKBP5 (ThermoFisher, Cat#711292, Lot 2311462, 1:200); guinea pig α-vGluT2 (EMP Millipore, Cat# AB2251-I, Lot 3593077, 1:1000); *Wisteria floribunda* lectin – fluorescein-labeled (Szabo-Scandic, Cat#VECFL-1351, Lot ZL0618, 1:200). The slices were then washed three times with PBS and incubated, protected from light, for 2 hours at room temperature on a shaker, with the secondary antibodies diluted in an antibody solution. The secondary antibodies raised in goat or donkey were purchased from Thermo Fisher Scientific (Alexa Fluor 488, Alexa Fluor 568, Alexa Fluor 647, 1:2000). The slices were washed three times with PBS. The nuclei were labeled with Hoechst 33342 (Thermo Fisher Scientific, Cat#H3570, 1:5000) diluted in PBS for 15 minutes. The slices were mounted on microscope glass slides (Assistant, Cat#42406020) with coverslips (Menzel-Glaser #0) using an antifade solution consisting of 10% (v/v) mowiol (Sigma, Cat#81381), 26% (v/v) glycerol (Sigma, Cat#G7757), 0.2M tris buffer pH 8, and 2.5% (w/v) Dabco (Sigma, Cat#D27802).

### Microscopy

#### Confocal microscopy

Images were acquired using Zeiss LSM800 or LSM900 upright microscopes with a Plan-Apochromat 40×/1.3 Oil DIC, UV-IR (Cat#420762-9800-799) or a Plan-Apochromat 40×/1.3 Oil DIC M27 (Cat#420762-9800-799), respectively. Z-stack images were obtained as a 2×2 tile scan, or 4×2 for whole VISp layers, at a resolution of 0.156×0.156×0.24 μm. Images from TRAP2 mice were captured with a Zeiss LSM800 using a Plan-Apochromat 20×/0.8 (Cat#420650-9901).

#### Super-resolution microscopy

The z-stack images of the dendritic spines in cortical layer II/III in Thy1-EGFP were acquired using a Zeiss LSM900 upright microscope equipped with a second-generation Airyscan module optimized for a Plan-Apochromat 40×/1.3 Oil DIC M27 (Cat#420762-9800-799). Images were acquired at a resolution of 0.045 × 0.045 × 0.19 μm.

### Image processing and analysis

Confocal tile images were converted to .ims using Imaris File Converter 9.9.1v and stitched with Imaris Stitcher 9.9.1.v. The images were processed using Imaris 9.9.1.v processing tool, applying a Gaussian filter width of 0.156 µm, setting the background subtraction to 76.2 µm, and activating the layers’ normalization.

#### Reconstruction of 3D segmented microglia

Microglial cells were traced in 3D with the filament-tracing plugin on Imaris 9.9.1v. Filament starting points were set on microglia soma with the largest diameter of 7.8 µm and seeding points of 0.5 µm. Disconnected segments were removed with a filtering smoothness of 0.5 µm. After the tracing, microglia cells at the image’s border with only partial tracing were manually removed. The data analysis was performed blinded. The vGluT2 staining, concentrated within layers I and IV (*69*), discriminated different cortical layers in the VISp. Microglia traces with somas located within layer I or IV were labeled as layer I and IV microglia, respectively, with layer II/III microglia in-between and layer V/VI microglia below. The generated layer-discriminated skeleton images were converted from .ims format (Imaris) to .swc format (*70*) by first obtaining the 3D positions and the diameter of each traced microglial process using the ImarisReader toolbox for MATLAB (https://github.com/PeterBeemiller/ImarisReader/) and then exporting for format standardization using the NL Morphology Converter (http://neuroland.org/). Artifacts from the 3D reconstructions automatically failed to be converted into .swc format.

#### Microglia morphological analysis with morphOMICs

3D segmented microglia were analyzed using morphOMICs, a topological data analysis (TDA) framework designed to extract and interpret microglial morphological phenotypes (Colombo et al., 2022). The morphOMICs pipeline comprises several sequential steps: First, the Topological Morphology Descriptor (TMD) (*71*) was applied using the radial distance from the soma as a filter function to capture the topological features of each segmented microglial structure via a persistence barcode. Then, persistence barcodes were filtered out based on the following criteria: (i) fewer than five or more than 250 bars; (ii) the longest bar exceeded 110 microns; (iii) more than 10 bars with a death distance of 0 microns. After visual inspection, we confirmed that these barcodes resulted from tracing errors, overlapping cells, wrongly segmenting two cells as one, or failures in the SWC file conversion. From our dataset of 3730 persistent barcodes, we excluded 7 persistence barcodes. Next, each barcode was vectorized using the persistence image transformation (*72*). Then, we applied multiple dimensionality reduction techniques to the persistence images to visualize microglia’s morphological embeddings and to explore condition-dependent morphological variation using Variational Autoencoder (VAE) (*73*). Before applying VAE, preliminary dimensionality reductions were performed. First, pixels exhibiting a standard deviation less than 1e-5 across all persistence images in the microglia dataset were removed. This step aimed to eliminate pixels consistently black across the dataset, typically located far from regions of high persistent feature density in the images. Subsequently, the dataset underwent z-score standardization to ensure each feature had a mean of zero and a standard deviation of one. Finally, Principal Component Analysis (PCA) (Pearson, 1901) was conducted using the implementation provided by scikit-learn with default parameters.

VAE was developed using PyTorch (v2.6.0; CUDA 12.6). The VAE encoder was used to capture the underlying data structure, and the decoder was used to reconstruct embedded persistence images for embedding (latent space) interpretation, respectively. Both were implemented as multilayer perceptrons. The encoder comprised three hidden layers with sizes [32, 16, 8], while the decoder mirrored the encoder architecture with layers of sizes [8, 16, 32]. Scaled Exponential Linear Unit (SELU) activation functions were employed for all hidden layers (*74*). The latent space was set to dimension 2 to facilitate visualization and interpretation. The VAE was trained on the first 64 principal components derived from the PCA applied to persistence images, which collectively accounted for over 99% of the total variance and facilitated efficient training. Weights within the network were initialized using Kaiming uniform initialization. The COntinuous COin Betting (COCOB) optimizer (*75*) was employed to eliminate the need for manual learning rate tuning. Training was conducted over 2,000 epochs with a batch size of 8. A Kullback-Leibler (KL) annealing strategy was implemented to balance the reconstruction loss and the KL divergence. Specifically, the weight of the KL term was gradually increased from approximately 0 to 1 following an exponential schedule, defined by the function *w*(*x*) = *1* − (*e* − *x*), where *x* values were linearly spaced between 2 and 7 (i.e. *x* = *5* ∗ *i*/*1999* + *2*, with *i* the epoch index between 0 and 1999). This approach enabled the model to focus on accurate reconstruction during the initial training phase, progressively encouraging the latent space to conform to a standard normal distribution.

An interpolation composed of five points was constructed by performing a linear regression in the latent space of the VAE on the medians of the female and male of both KXA and Saline to compute the slope and intercept. The 2D embedded persistence images in the VAE latent space were then projected onto this interpolation line, resulting in a one-dimensional representation of the distribution for the four groups. After assessing normality using the Shapiro-Wilk test, we performed a Kruskal–Wallis test, followed by a post hoc Dunn’s test with Bonferroni correction for multiple comparisons. Statistical significance was determined at a p-value threshold of 0.05. A Jupyter notebook reproducing the results in this paper and the corresponding parameter file can be found at https://github.com/siegert-lab/V1_morphOMICs.

#### CD68 volume within microglia

Surface renderings were generated on microglia and CD68 z-stacks using the surface-rendering module of Imaris 9.9.1.v. Surfaces were created with the surface detail set to 0.2 µm. The surface-surface coloc plugin was utilised to identify the CD68 surface within microglia. This analysis was performed on the entire image. The total ratio of CD68 volume to microglial volume (CD68-to-microglial volume) was calculated for each image.

#### PNN inside CD68 volume within microglia

Surface renderings of microglia, CD68, and PNN were generated from z-stacks using the surface-rendering module of Imaris 9.9.1.v. Surfaces were created with a surface detail setting of 0.2 µm. The CD68 surface within microglia was first identified using the surface-surface colocalization plugin, which generated a new surface module. Subsequently, the surface-surface colocalization plugin was applied to identify overlapping surfaces between the microglia-CD68 colocalization and the PNN. The ratio of PNN within the microglial CD68 volume was calculated for each image.

#### Dendritic spines quantification

The density of dendritic spines from super-resolution images was quantified semi-automatically using Imaris 9.9.1v. The manual drawing module was activated through the filament tracing function. A starting point was established at the beginning of the dendrite, and the dendrite was traced with the auto-path function. The spine density was automatically calculated using the Imaris filament creation module by entering the dendrite diameter, the thinnest point (corresponding to the spine neck), and the maximum length measured for each dendrite.

### Slice electrophysiology recording

Acute parasagittal brain slices containing the primary visual cortex (VISp) were derived from P60-P90 C57BL/6J male and female mice. Four hours after the injection of KXA or saline, the mice were anesthetized with isoflurane and decapitated. Following a midline sagittal incision, the brain was rapidly removed, mounted, and 250 µm thick parasagittal slices were cut on a vibratome (VT1200S, Leica) in ice-cold cutting solution (pH 7.3, adjusted with HCl) containing (in mM): 135 N-methyl-d-glucamine, 1 KCl, 1.2 KH2PO4, 10 glucose, 20 choline bicarbonate, 1.5 MgCl_2_, and 0.5 CaCl_2_, and continuously oxygenated with 95% O_2_/5% CO_2_. The acute brain slices were incubated at 32°C in oxygenated artificial cerebrospinal fluid (ACSF) containing (in mM): 118 NaCl, 2.5 KCl, 1.25 NaH_2_PO_4_, 2 MgCl_2_, 2 CaCl_2_, 26 NaHCO_3_, 3 Myo-Inositol, 20 Sucrose and 10 glucose for 30 min, followed by a slow cooldown to RT over an additional 30 min. For recordings, brain slices were transferred to a recording chamber on an LNScope 240 XY (Luigs & Neumann) and superfused with ACSF (22-24 °C) at a rate of 3 mL min^−1^ with a peristaltic pump (MP3, Gilson). Layer II-III pyramidal neurons were recorded through patch pipettes (3–5 MΩ) made from borosilicate glass capillaries pulled on a P1000 glass puller (Sutter Instruments). Patch pipettes were filled with an intracellular solution containing (pH 7.3, adjusted with KOH; 302 mOsm; in mM): 120 K-gluconate, 20 KCl, 0.5 EGTA, 2 MgCl_2_, 10 HEPES, 2 Na-ATP, 0.2 Na-GTP, 23 Sucrose, and 0.5% biocytin (w/v) (Tocris, 3349). mEPSCs were recorded in voltage-clamp at -60 mV, in the presence of TTX 1 µM (Tocris, 1069) and Bicuculline 10 µM (Tocris, 0109), using a Multiclamp 700B amplifier (Molecular Devices) connected to a 1550B digitizer (Molecular Devices), sampled at 10 KHz, and filtered at 2 KHz. The frequency and amplitude of the mEPSCs were analyzed using the Mini Analysis Program (Synaptosoft).

### *In-vivo* live imaging

#### Cranial window surgery

Cx3Cr1Cre^ERT2/het^×Ai9×Thy1-GFP mice, 8-10 weeks old, were anesthetized with isoflurane (Zoetis, Cat#6089373, 5% induction, 2.5% maintenance) in 0.6 L/min O_2_ and were maintained at 37°C using a heating pad connected to a rectal probe during the surgical procedure (Harvard Apparatus, Cat#14-516-264). The mice were positioned in a stereotaxic frame (KOPF digital plus).

Eye ointment (Oleovital) was applied to prevent corneal dehydration. The head was shaved and disinfected with 70% (v/v) EtOH in H_2_O; an incision in the scalp was made, and the skin flap covering both hemispheres was removed. The periosteum was treated with a 3% (v/v) hydrogen peroxide (Sigma, Cat#216763) in PBS solution. A 3 mm biopsy tool (Henry Schein Cat#394-314) was used to mark the skull above VISp for drilling, and a craniotomy centered 3.6 mm posterior to Bregma and 1.65 mm interaural was performed on the right hemisphere with a 0.9 mm micro drill (Fine Science Tools Cat#19007-09). After achieving hemostasis, the exposed brain was covered with a double glass window made of 3- and 5-mm glass coverslips (Multi Channels System Cat#640720 and Cat#640731) glued together with glass superglue (UHU Glass Special Glue). The glass window was then fixed to the skull using dental cement (Kulzer Paladur Cat#64707948 and Cat#64707938). A custom-made metal frame was secured to the skull with dental cement. Pain control was administered with Metamizol (Sanofi Aventis, Cat#Ay005, *s.c.* 200 mg/Kg during surgery) and Meloxicam (Boehringer-Ingelheim, Cat#KPOEH3R, *s.c.* 5 mg/Kg after surgery every 24 h for three consecutive days). The animals recovered for at least two weeks before being used for experiments.

#### 2-photon microscopy

Awake Cx3Cr1^CreERT2/het^×Ai9×Thy1-GFP mice were head-fixed on a custom-made microscopy stage and imaged with a Leica TCS SP8 DIVE CS microscope equipped with an HC FUOTAR L 25×/0.95 W (Cat#15506374), WD=2.5 mm, wide angle (41°). Two lasers were used in this experiment: the tunable laser set at 920 nm and the non-tunable one at 1045 nm. Approximately 100 µm z-stack images at a resolution of 1600×1600 pixels and a z-step of 2 µm were acquired starting 50 µm below the dura. The attenuation correction was activated on the laser power to compensate for signal reduction during the z-stack acquisition. Images were acquired bidirectionally with an x-phase of 31.55 using the two HyD SP GaAsP-Detectors, one for GFP and one for Ai9, with a set gain of 100. The time interval between the z-stacks was 5 minutes. The experiment began with 30 minutes of awake recording. Imaging was paused, and the animal received KXA or KXA+corticosterone (20 mg/Kg) *i.p.* injection. Imaging was then resumed, reaching a total duration of 210 minutes.

#### Image processing

2-photon timelapse images were converted to .ims using Imaris File Converter 9.9.1v. A spot with a diameter of 8 µm was created on the soma of each microglia cell using the Imaris spot function. A translational drift correction of image and spot objects was performed. Images were processed with Imaris 9.9.1.v processing tool, utilizing a Gaussian filter width of 0.174 µm, a background subtraction set to 11.2 µm, and activated normalization of the layers.

### Single-nucleus multiome sequencing

#### Tissue preparation

C57BL/6J females, eight weeks old, littermates in the follicular phase, received KXA or saline *i.p.* injection. Two hours after the treatment, the animals were anesthetized with isoflurane and perfused with ice-cold HEPES cutting solution containing (in mM): 110 NaCl, 10 HEPES, 25 glucose, 75 sucrose, 6 MgCl_2_, 7.5 MgCl_2_, and 2.5 KCl. The pH of the solution was adjusted to 7.4 using 1N NaOH. After being perfused with 20 mL of ice-cold HEPES cutting solution, the brain was explanted and immediately frozen in liquid N_2_ vapors for 2 minutes. The brains were stored at -70°C in 5 mL Eppendorf tubes containing OCT (A. Hartenstein, Cat#TTEK). The samples were shipped to the Allen Institute for Brain Science.

#### Sequencing

##### cDNA amplification and library construction

For 10x Multiome library generation, we utilized the Chromium Next GEM Single Cell Multiome ATAC + Gene Expression Reagent Bundle (10× Genomics, Cat#1000283). We adhered to the manufacturer’s instructions for transposition, nucleus capture, barcoding, reverse transcription, cDNA amplification, and library construction (Allen Institute for Brain Science. 10x Multiome sample processing. Protocols.io https://doi.org/10.17504/protocols.io.bp2l61mqrvqe/v1 (2023)).

For the snMultiome libraries, we loaded 11,154 ± 1,386 nuclei per port. For snRNA-seq, we targeted a sequencing depth of 120,000 reads per nucleus. The actual average achieved for the nuclei included in this study was 87,183 ± 24,943 reads per nucleus. For snATAC-seq, we targeted a sequencing depth of 85,000 reads per nucleus. The actual average achieved for the nuclei included in this study was 90,470 ± 7,780 reads per nucleus across 4 libraries. The snMultiome libraries were sequenced on the Illumina NovaSeq6000, and sequencing reads were aligned to the mouse references downloaded from 10× Genomics, which includes ensembl GRCm38 (v98) fasta and gencode (vM23) gtf file, using the 10× Genomics CellRanger Arc (v2.0) workflow with default parameters.

### Data analysis

#### Quality control

We loaded the UMI count matrices for each mouse sample in R (v4.3.1, Linux distribution) as a Seurat object (*76*), where each gene must be expressed in at least five captured nuclei, and each cell must contain at least 300 genes with non-zero counts. For each sample, we implemented scDblFinder (*77*) to detect potential doublets—two nuclei that were sequenced as one—with the following parameters: “min.mean” : 0.1, “includePCs” : 20, “nfeatures” : 1250, and “seed” : 34151 for reproducibility. After filtering out the inferred doublets, we further filtered out nuclei with total counts and unique genes greater than 3MAD (median absolute deviation) across all nuclei sequenced per sample (**Figure S5A-B**). To focus more on canonical genes and further reduce noise, we drop pseudogenes or non-coding RNAs, i.e., genes whose names start with “Gm” or end with “Rik”, as well as microRNAs (genes starting with “Mir” and ending with “hg”).

#### Normalization and integration

We log-normalized the filtered count matrix using Seurat’s NormalizeData(). We identified 3000 highly variable genes with the vst selection method using FindVariableFeatures(), which we used as anchors to integrate the samples with SelectIntegrationFeatures(), FindIntegrationAnchors(), and finally, IntegrateData(). We scaled the integrated dataset using ScaleData() and conducted an initial PC reduction using RunPCA(), taking only the first 50 PCs. To visualize potential batch effects, we inferred a 2D manifold using RunUMAP() with the following parameters: “umap.method” : “umap-learn”, “metric” : “cosine”, “n.neighbors” : 100, “n.components” : 2, “spread” : 1, “densmap” : 0, and “seed.use” : 5143 for reproducibility (**Figure S5C-D**).

#### Cell type assignment

To extract high-resolution clusters of nuclei with similar transcriptomic profiles, we implemented the Louvain algorithm using FindNeighbors() with the following parameters: “k.param” : 20, “nn.method” : “annoy”, “annoy.metric” : “cosine”, “l2.norm” : 0, and then with FindClusters() with the following parameters: “resolution” : 0.5, “algorithm” : 1 for the original Louvain algorithm, and “random.seed” : 3617 for reproducibility. We then used SingleR (*78*) to annotate cluster identities with the Allen Brain Atlas VISp transcriptomic cell type annotations (*79*) as a reference. Note that this Allen Brain annotation contains cellular subtypes based on specific gene markers. In this paper, we focus on the main cell type annotations. We confirm the cluster identities using known gene markers (**Figure S5E-H**).

#### Interaction model

We estimated the interaction effect of ketamine on neuronal versus astrocytes/microglia cell types while accounting for variability across mice using a linear mixed model (LMM). For fixed effects in the LMM, we estimated the ketamine effect, the cell type effect, and the interaction between ketamine and cell type. We accounted for variability across mice as random effects. We fit the model for each of the top 2000 highly variable genes, calculated using binomial deviance (*80*).

For each gene *g*, we fit a linear mixed model:

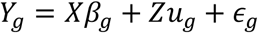

Where:

*Y*_*g*_ is the log-normalized mRNA abundance for gene *g*. We calculated *Y*_*g*_ by log transforming and normalizing the raw counts 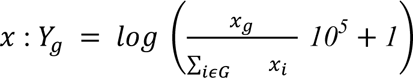 variable genes.

*X* is the design matrix for fixed effects. For the full model, it is N by 4, and for the null model, it is N by 3 (N=37,054 cells). The four columns denote (1) intercept, (2) whether the cell is an astrocyte/microglia cell type, (3) whether the cell is treated with ketamine, or (4) whether the cell is both an astrocyte/microglia cell type and treated with ketamine. In this design matrix, for example, a neuronal cell type without ketamine treatment (intercept case) would be (1, 0, 0, 0). In the null model, we dropped the fourth column to assume no interaction.

*β* is a vector of fixed effects, which are log fold changes for different conditions. For the full model, the four fixed effects are (1) intercept (log mRNA abundance of a neuronal cell type with no ketamine treatment), (2) log fold change for being an astrocyte/microglia cell type, (3) log fold change of ketamine treatment, or (4) interaction effect of ketamine treatment and astrocyte/microglia cell type.

*Z* is the design matrix for random effects of size N by m, where m is the number of mice in the experiment (m=4 mice).

*u* is a vector of random effects.

We fit the data with the full model and compared it with the null model to find genes where the interaction between ketamine treatment and astrocytes/microglia cell type was significant. We fit the full model using the lme4 R package with the R formula:

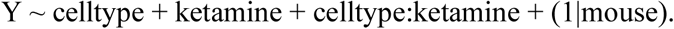

For the null model, we set the interaction effect to zero:

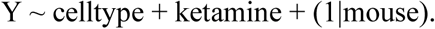

We compared the two model fits using ANOVA implemented in lme4 to obtain a p-value: anova(null_model, full_model)

which uses an F-statistic calculation (*81*).

### Fluorescence *in situ* hybridization

C57BL/6J mice of both sexes, aged 8–10 weeks, were perfused with 4% (w/v) PFA in PBS two hours after the injection of KXA or saline, in line with the timeline of the single-nucleus multiome sequencing experiment. The explanted brains were post-fixed in 4% (w/v) PFA in PBS for 24 hours at 4°C on an orbital shaker. Tissues were cryoprotected in an ascending gradient of sucrose (10% (w/v), 20% (w/v), and 30% (w/v) in PBS) until the tissues sank to the bottom of the container (approximately 18 hours per step). The brains were embedded in OCT (A. Hartenstein, Cat#TTEK), and 10 µm slices were cut using a cryostat (Fisher Scientific, NX70, Cat# 957000L). The slices were stored at -70°C. The fluorescence *in situ* hybridization experiment was conducted using the RNAScope Intro Pack for Multiplex Fluorescence Reagent Kit v2 (Bio-techne, Cat# 323136), following the protocol provided by the manufacturer. In this experiment, we identified the following probes: FKBP5 (Probe - Mm-Fkbp5 – Mus musculus FK506 binding protein 5 (Fkbp5) mRNA, Cat#457241), Cx3cr1 (Mus musculus chemokine (C- X3-C) receptor 1 (Cx3cr1) mRNA, Cat#314221-C2), Rbfox3/NeuN (Probe - Mm-Rbfox3-C2 - Mus musculus RNA binding protein fox-1 homolog (C. *elegans*) 3 (Rbfox3/NeuN) transcript variant 1 mRNA, Cat#313311-C2), and S100β (Probe - Mm-S100b-C3 - Mus musculus S100 protein beta polypeptide neural (S100b) mRNA, Cat#431731-C3). The probes were visualized with TSA Vivid Dyes linked to 520, 570, or 650 nm fluorophores (Bio-techne, Cat#323271, 323272, and 323273, respectively). The nuclei were labeled using DAPI (available in the kit). The z-stack images were acquired in super-resolution using a Zeiss LSM 900 upright microscope, equipped with an Airyscan 2^nd^ generation module optimized for a Plan-Apochromat 40×/1.3 Oil DIC M27 (Cat#420762-9800-799). Images were captured at a resolution of 0.045×0.045×0.19 μm. The images were analyzed in Imaris 9.9.1v using the Spot function. The results were expressed as puncta density (n puncta/area).

### Blood corticosterone measurement

C57BL/6J mice of both sexes, 8 to 10 weeks old, were injected with saline or KXA. Thirty minutes or two hours after the injection, the mice were anesthetized with isoflurane, and the skin was sterilized using 70% (v/v) ethanol before the chest and abdomen were opened. The *inferior vena cava* was cut, and the systemic blood was collected in a 2 mL Eppendorf tube. The blood was allowed to coagulate for 15 minutes at room temperature. The samples were centrifuged (Eppendorf 5224R cooling, Cat# 5424000410) at 4°C for 10 minutes at 2000×g to separate the serum. The supernatant was collected and stored at -70°C. The corticosterone was quantified using an ELISA competitive kit (Biocat, Cat#K014-C1-AAS) following the manufacturer’s instructions. The results were read with an ELISA plate reader (Biotek SynergyH1).

### Adrenalectomy surgery

C57BL/6J females, 8 to 10 weeks old, were anesthetized with isoflurane (5% induction, 2.5% maintenance) at a rate of 0.6 L/min O_2_ and were maintained at 37°C using a heating pad connected to a rectal probe during the surgical procedure. After shaving and sterilizing the skin, adrenalectomy was performed using a dorsal midline incision approximately 1 cm long, located at the level of the third lumbar vertebrae, *caudal* to the last rib, to access the abdominal cavity. The kidney, adrenal gland, and associated fat pad were identified, and the vessels supplying the gland were cauterized. The gland was then excised at its base. The procedure was performed bilaterally. The abdominal cavity and skin were sutured using Dafilon Blue 4/0 1.5 surgical sutures (Henry Schein, Cat#656-476). To prevent changes in circadian rhythm, adrenalectomized mice were given 25 µg/mL corticosterone in drinking water, approximating basal corticosterone levels and mimicking the typical circadian secretion pattern (*51*). The mice were allowed to recover for at least two weeks. Pain was managed with Metamizol (Sanofi Aventis, Cat#Ay005, *s.c.* 200 mg/Kg during surgery) and with Meloxicam (Boehringer-Ingelheim, Cat#KPOEH3R, *s.c.* 5 mg/Kg after surgery every 24 hours for three consecutive days.

### Statistics

Data were analyzed using GraphPad Prism 10.2.2. The datasets were tested for normal distribution using the Shapiro-Wilk test. For multiple comparison analyses of normally distributed data, we used a two-way ANOVA. If a significant effect (p < 0.05) was found, we corrected for multiple comparisons using Tukey’s correction. We used a Friedman test for multiple comparison analyses of non-normally distributed data. Two sets of unpaired variables that showed normal distribution were compared using Student’s t-test, whereas, in the case of non-normal distribution, the Mann-Whitney test was used. Data in the plots are expressed as means ± standard error of the mean. Statistical significance across groups was determined using the non-parametric Kruskal-Wallis test. If a significant effect (p < 0.05) was found, post hoc pairwise comparisons were conducted using Dunn’s test to account for multiple comparisons. Group differences with unequal variances across groups were analyzed using Welch’s ANOVA test. When a significant effect was detected, Dunnett’s T3 multiple comparisons test was used to compare each treatment group to the control group. Statistical significance was indicated as \**p* < 0.05, \*\**p* < 0.01, or \*\*\**p* < 0.001. ^ns^*p* > 0.05, not significant.

If not otherwise indicated, graphs show mean and standard error of the mean (SEM). A detailed statistical analysis is available in the **Supplementary Table**.

## Code availability

Codes are available here for morphological analysis: https://github.com/siegert-lab/morphOMICs_Venturino_2025

Single-nucleus multiome analysis: https://github.com/jakeyeung/GliaMultiomics

## Data availability

Transcriptional data deposit under GSE298669.

## Author contributions

Conceptualization and experimental design: A.V., S.S.; Investigation, validation, and methodology: A.V., M.A., for electrophysiology P.K., for multiome K.J., C.T.J.V., B.T.; Formal Analysis and software: T.N. and M.A. (morphOMICs), R.J.A.C (quality control of Multiome), J.Y. (KXA estimates); Writing – Original Draft: S.S.; Visualization: A.V., S.S.; Writing – Review & Editing, A.V., B.T., S.S.; Supervision and funding acquisition: S.S. All authors reviewed and approved the final manuscript.

## Acknowledgement

We thank the scientific service units at ISTA, especially the imaging optic facility (IOF), the machine shop (Miba), specifically Todor Asenov, the preclinical facility (PCF), specifically Michael Schunn and Sonja Haslinger, Christian Jansen for bioinformatic support, Marco Benevento for supporting the design and partial performance of electrophysiology and fluorescence *in situ* hybridization experiments, and for extensive discussions about the manuscript draft; Rouven Schulz and Gloria Colombo, for advice on morphological reconstruction; Mathias Schmidt for valuable discussions about Fkbp5; and all Siegert team members for constant feedback on the project and manuscript. Grammarly was used to assist with grammatical revisions.

## Figure legend

**Supplementary Figure 1.**
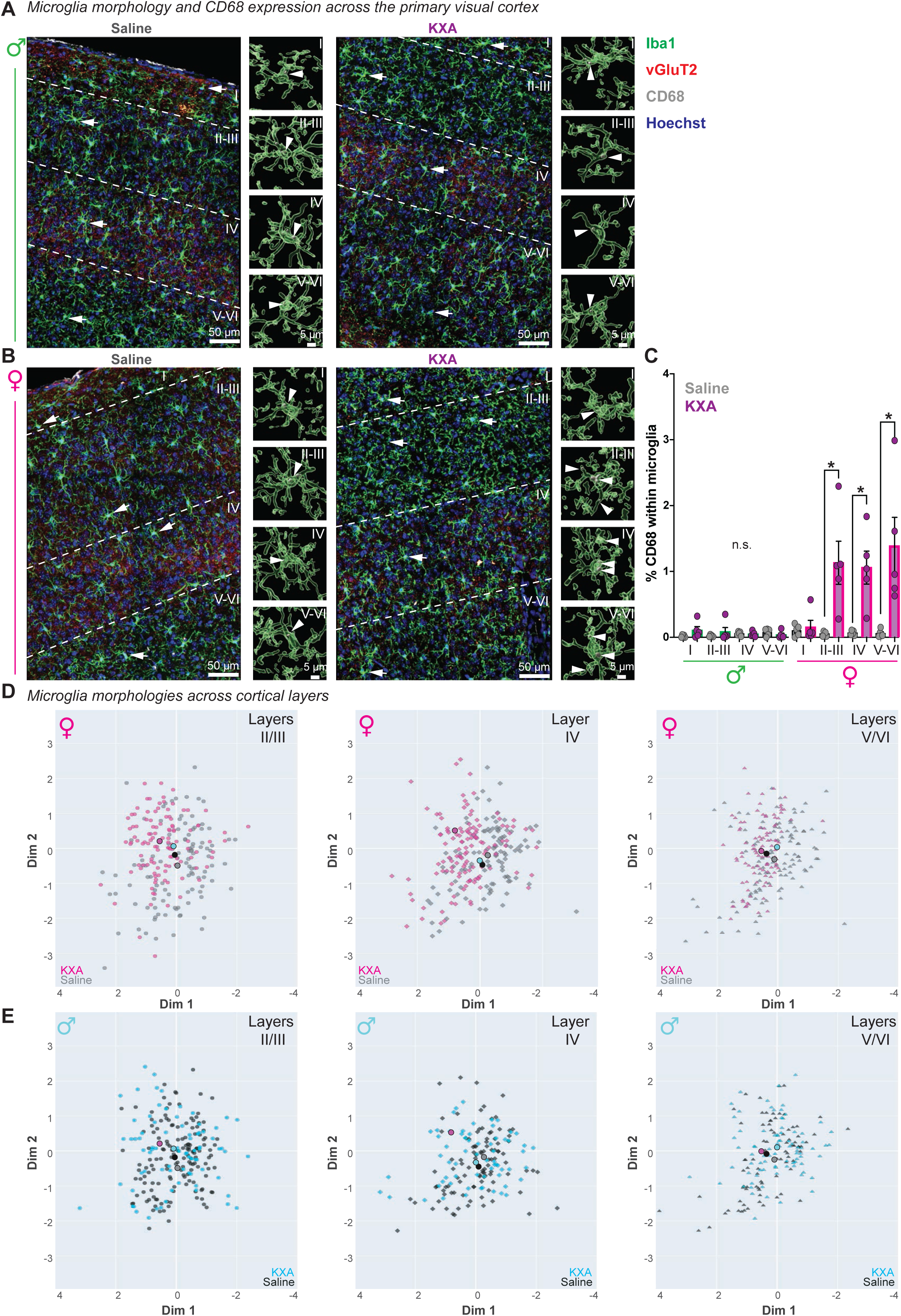
CD68 expression across VISp cortical layers and microglia morphological adjustment 4h after ketamine anesthesia in males and females. (**A-C**) CD68 quantification across the primary visual cortex (VISp) of males (**A**) and females (**B**), 4 hours after saline or KXA (ketamine-xylazine-acepromacine) injection. (**A-B**) Representative immunostainings for Iba1 (green), CD68 (magenta), vGluT2 (red), counterstained with the nuclei-dye Hoechst (blue). Left, overview image of the cortical layers assigned based on the vGluT2 signal. Arrow in each layer, microglia chosen for 3D surface rendering, shown next to the overview. Arrowhead, CD68 location within microglia. Scale bars: 50 µm for overview and 5 µm for rendering. (**C**) Bar chart of the mean percentage of CD68 volume within microglia in each cortical layer with SEM. Each dot, one animal. 5 animals/condition. Kruskal-Wallis with Dunn’s multiple comparison post hoc test, *p < 0.05, ^ns^*p* > 0.05, not significant. (**D-E**) Morphological analysis of female (**D**) and male (**E**) microglia within cortical layer II/III, IV, and V/VI in VISp after saline (grey, black) or KXA (magenta, cyan) using morphOMICs. Each microglia’s persistence image was embedded in a latent space using a Variational Autoencoder (VAE, see Methods). Larger dots, population mean for sex and condition.

**Supplementary Figure 2.**
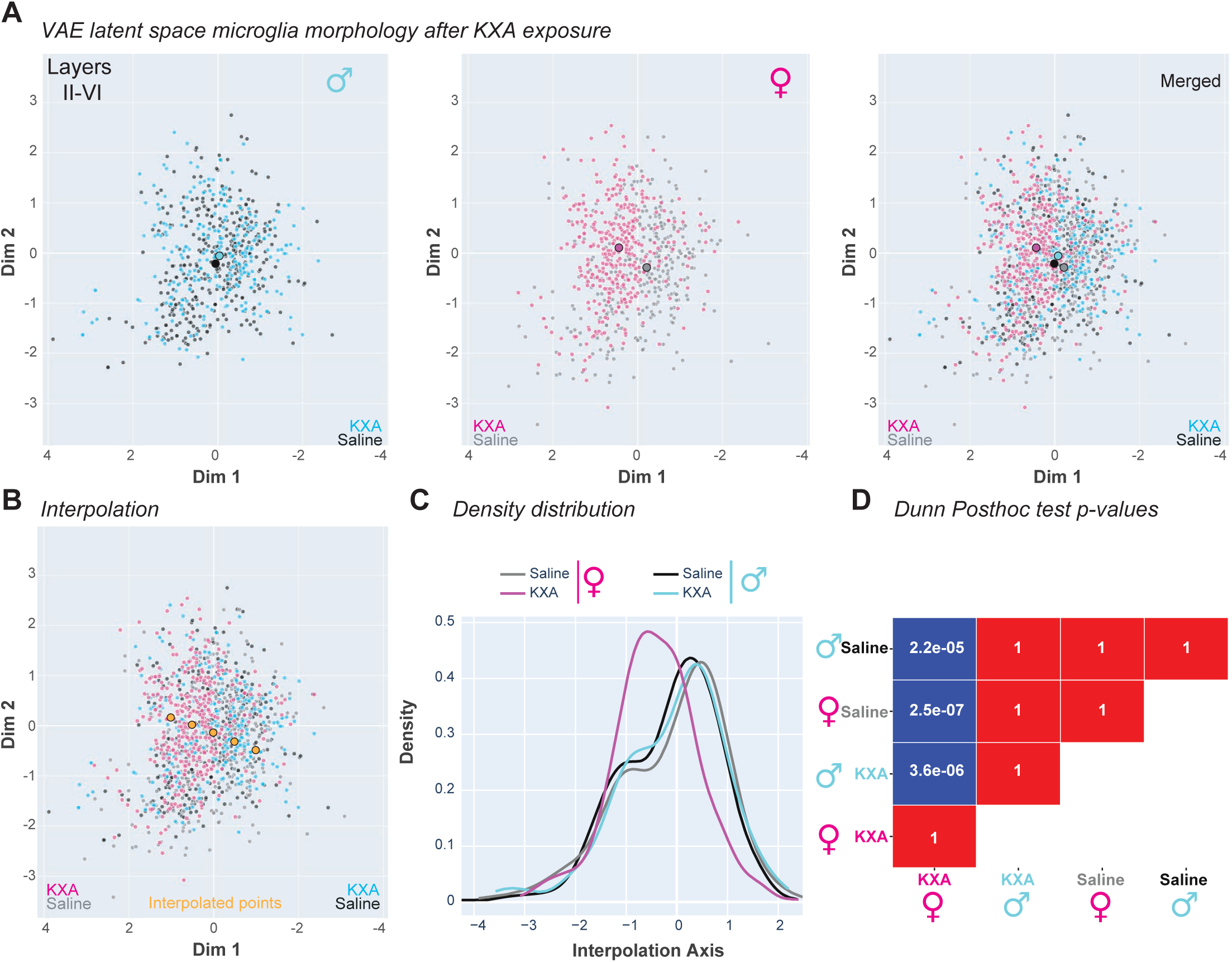
Sex-specific morphological differences in microglia 4h after ketamine anesthesia. (**A-D**) Morphological comparison between male (cyan) and female (magenta) microglia across the cortical layers II-VI in the primary visual cortex (VISp) after saline (grey, black) or KXA (magenta, cyan) using morphOMICs. (**A**) Variational Autoencoder (VAE) latent space representation. Each dot, a single persistence image. Larger dots, population median for sex and condition, color-coded by group conditions. (**B**) One-dimensional projection of the density distribution of embedded persistence images along the interpolation axis. (**C**) Density distribution computed by projecting the embedded persistence images along the interpolation axis (**B**), giving a one-dimensional projection. Color-coded by group conditions. The distribution shift highlights the distinctiveness of the female KXA group relative to the others. (**D**) Statistical comparison of microglia populations using Shapiro-Wilk with Dunn’s post hoc test with p-values.

**Supplementary Figure 3.**
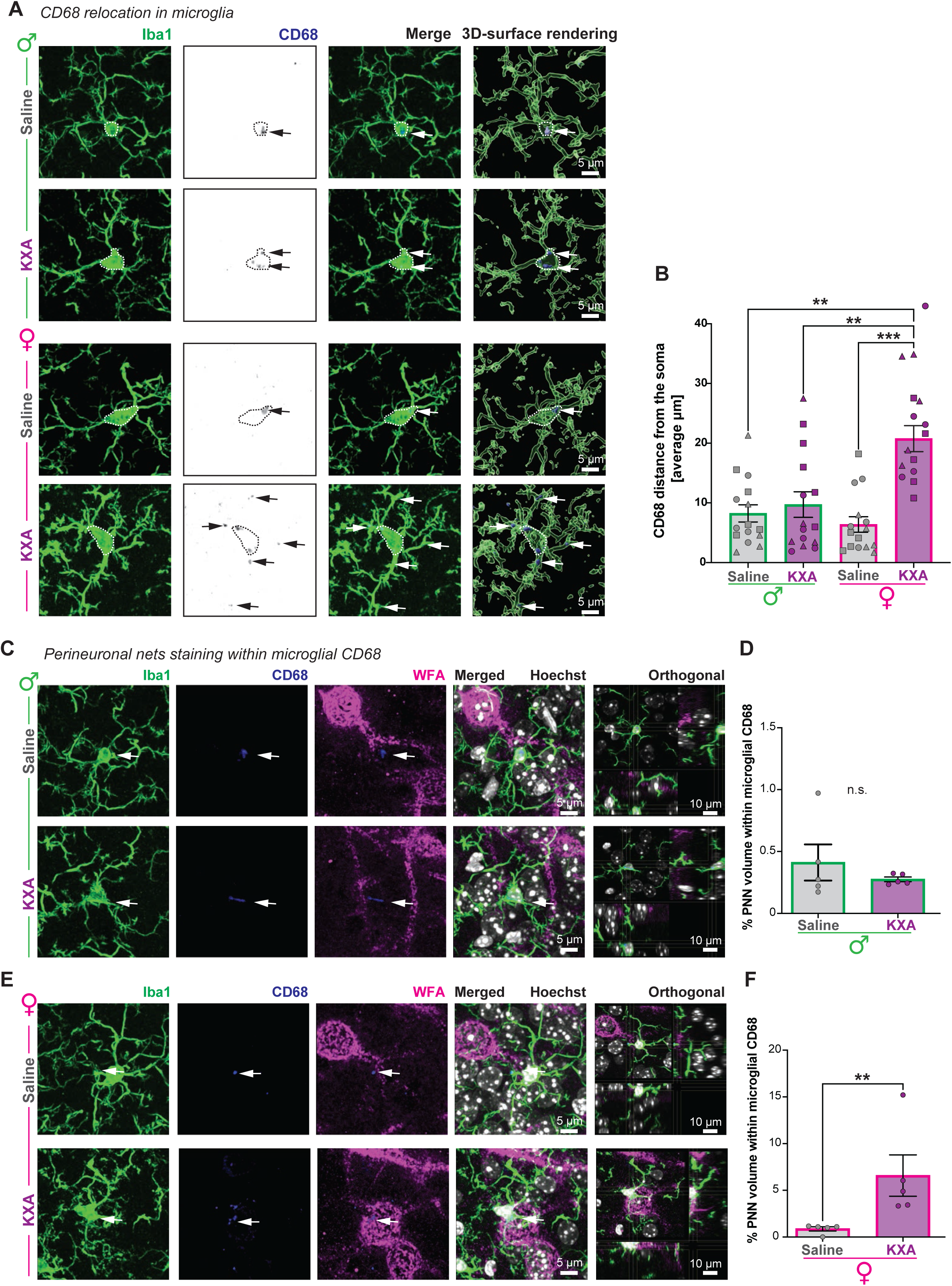
KXA stimulates female microglia to relocate CD68 and to engulf perineuronal nets (PNN) fragments. (**A-F**) Immunostainings for microglia with Iba1 (green) and CD68 (blue) in the primary visual cortex (VISp), layer III-V of males and females, 4 hours after saline or KXA (ketamine-xylazine-acepromazine) injection. (**A-B**) Comparison of CD68 relocation within microglia between males (green) and females (magenta). (**A**) Representative immunostainings across sex and condition. Dashed circle, contour of microglia soma as reference. Arrow, CD68 location within microglia. Next to merged images, 3D-surface rendering. Scale bar: 5 µm. (**B**) Bar chart of average CD68 distance from the microglia soma with SEM. Each dot, a cell with 5 cells/animal. Each symbol, one animal with 3 animals/condition. Kruskal-Wallis with Dunn’s multiple comparison post hoc test, **p < 0.01, ***p < 0.001. (**C-F**) Comparison of perineuronal nets staining inside microglia CD68 between males (green, **C-D**) and females (magenta, **E-F**). (**C**, **E**) Representative immunostainings across sex and condition, including *Wisteria floribunda agglutinin* (WFA, magenta) staining for perineuronal nets (PNN), and counterstained with the nuclei-dye Hoechst (white). Arrow, CD68 inside microglia. Scale bars: 5 µm and 10 µm for the orthogonal projection. (**D**, **F**) Bar charts of the mean percentage of PNN volume within microglial CD68. Each dot, one animal. 5 animals/condition. (**D**) Unpaired t-test with Welch’s correction. ^ns^*p* > 0.05, not significant. (**F**) Mann-Whitney test, **p < 0.01.

**Supplementary Figure 4.**
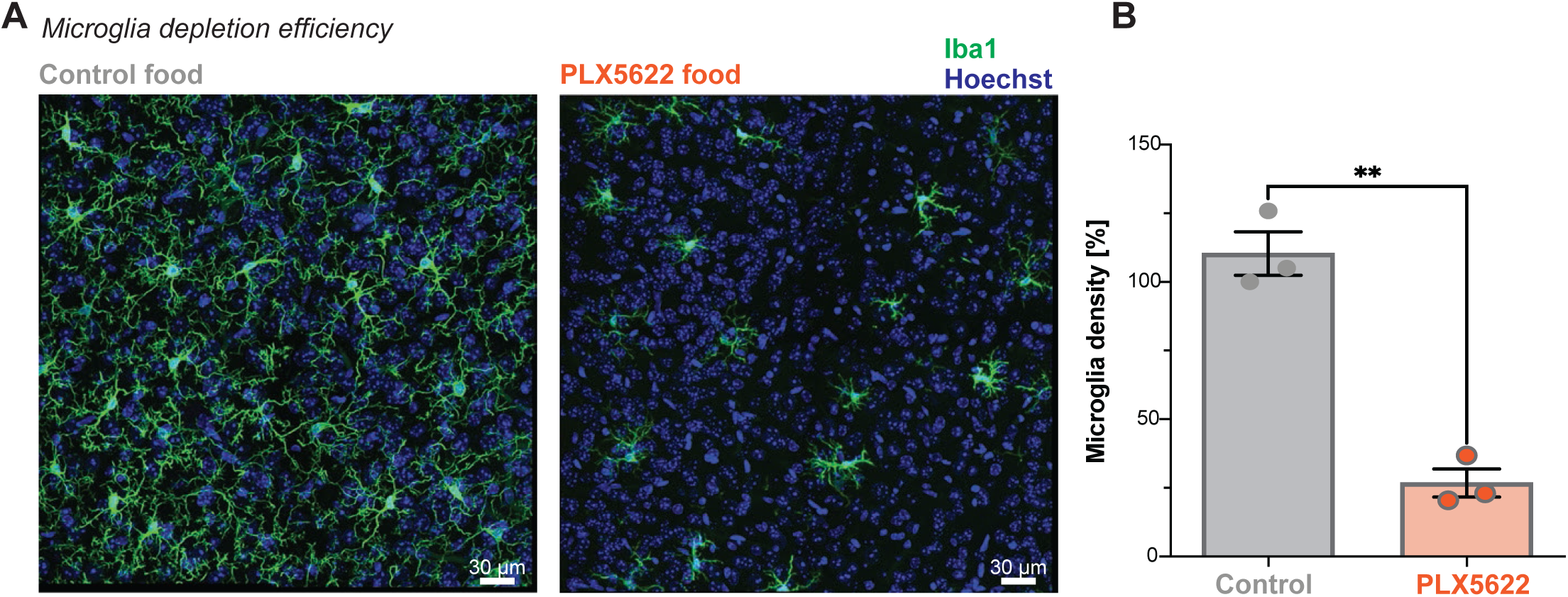
Microglia depletion efficiency with PLX5622. (**A**) Representative immunostainings for Iba1 (green), counterstained with the nuclei-dye Hoechst (blue). Left, control food. Right, feeding animals for 1.5 weeks with chow containing the Csf1-receptor inhibitor PLX5622. Scale bar: 30 µm. (**B**) Bar chart of mean percentage of microglia density in the VISp of saline and PLX5622-treated mice with SEM. Each dot, one animal. 3 animals/condition. Unpaired t-test with Welch’s correction, **p < 0.01.

**Supplementary Figure 5.**
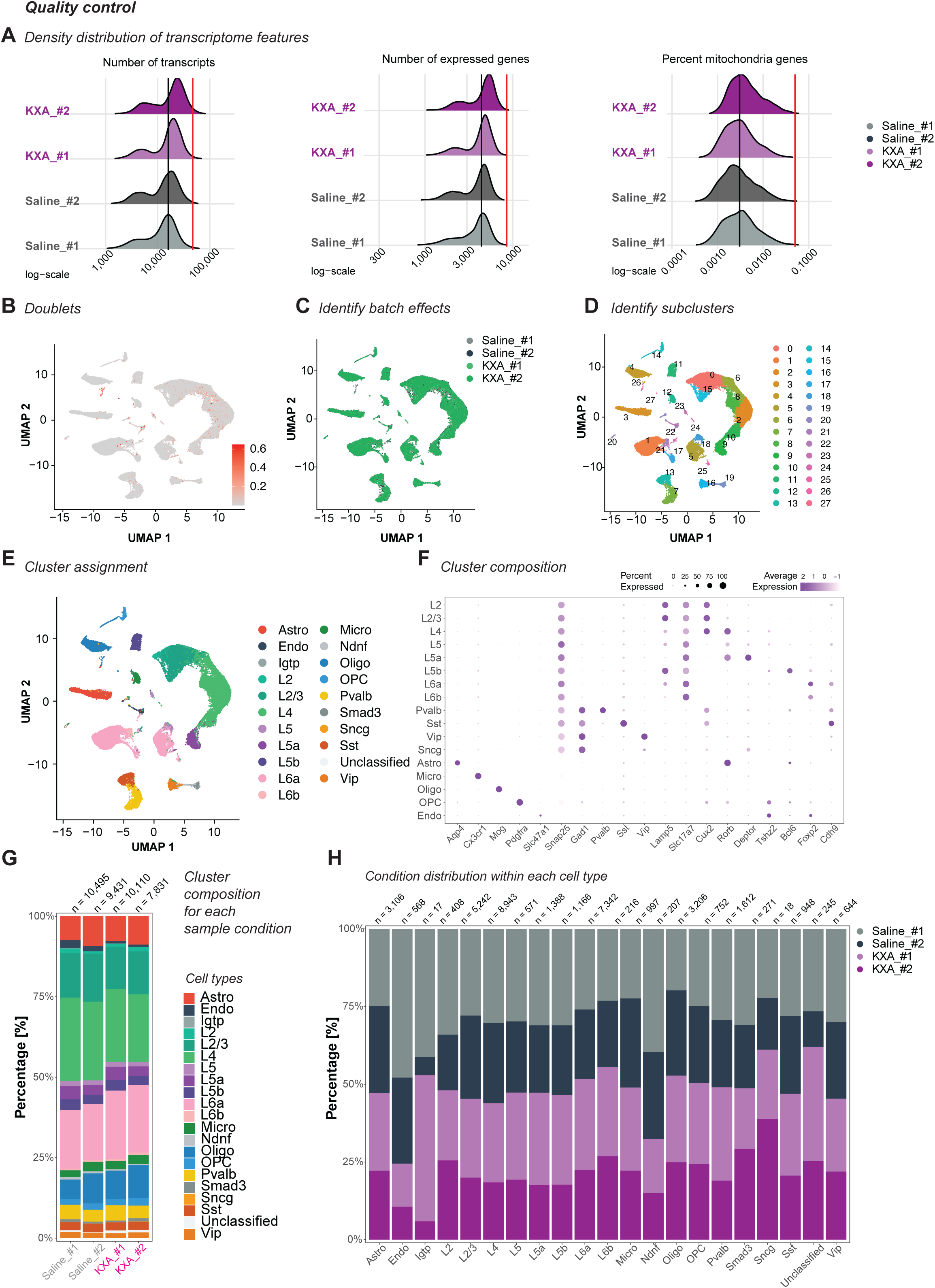
Quality control of single-nuclei multiome sequencing data. (**A-H**) Multiome single-nuclei sequencing data from the entire primary visual cortex (VISp) of females, 2 hours after saline or KXA (ketamine-xylazine-acepromazine) injection. For each condition, two animals. Detailed description in the method section. (**A**) Density distribution plots of transcriptomic features focusing on the total number of transcripts (left), the number of expressed genes (middle), and the percentage of mitochondrial gene expression (right). Black line, median. Red line, threshold for filtering. (**B**) UMAP representation for doublets within the dataset. Doublets were filtered out. (**C**) UMAP plot after log-normalization of the filtered count matrix. (**D-E**) UMAP of batch corrected representation and assignment of subclusters to cell types. (**F**) Verification of cell-type-specific signature genes in the corresponding cluster. (**G-H**) Distribution plots for the percentage of cell type populations within a condition (**G**) and the percentage of conditions within a cell type population (**H**). Absolute cell numbers are indicated above each bar.

**Supplementary Figure 6.**
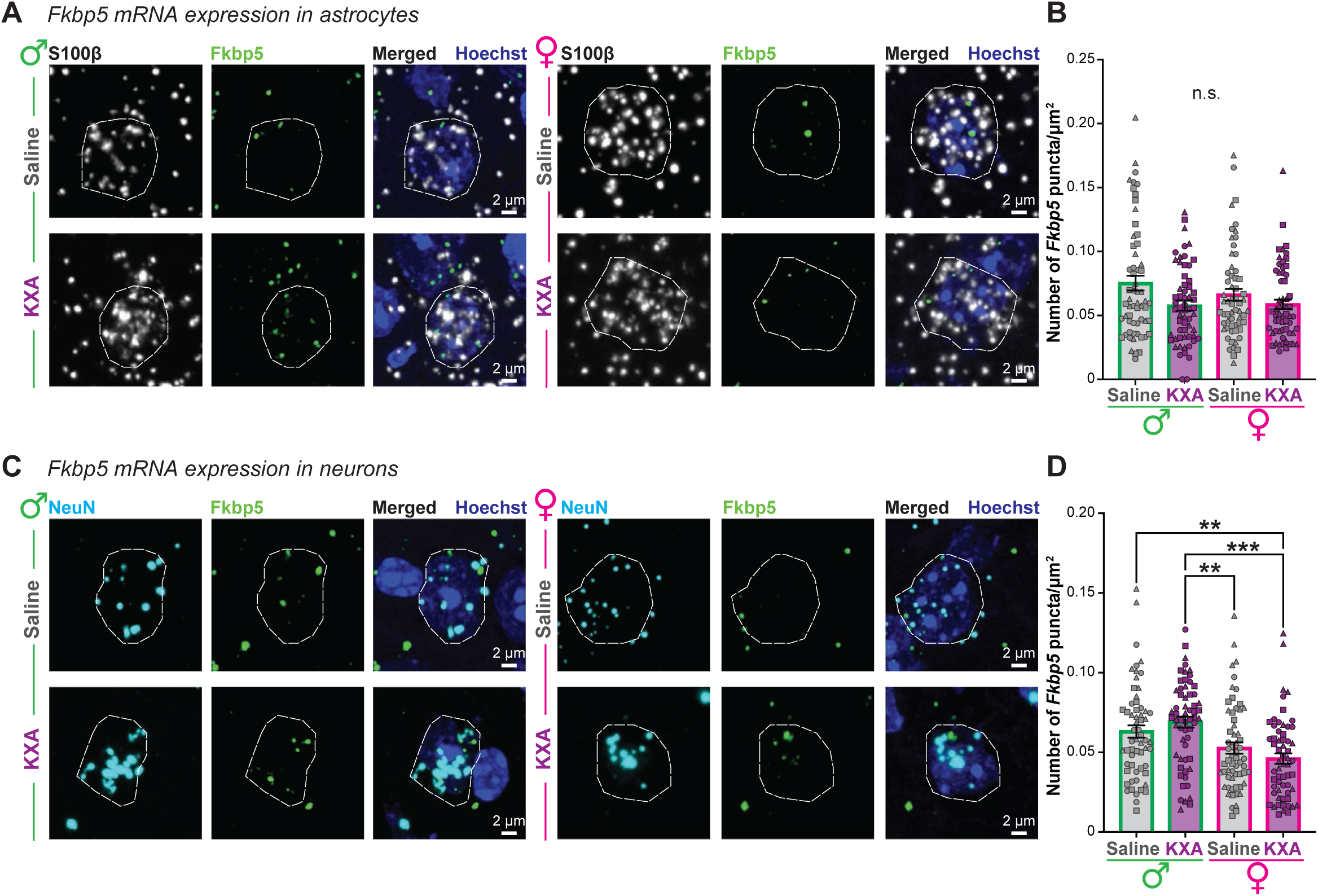
Astrocytes and neurons do not upregulate *Fkbp5* mRNA expression during ketamine anesthesia recovery. (**A-D**) Fluorescence *in situ* hybridization for male and female mice 2 hours after saline or KXA injection in VISp for mRNA probes against *Fkbp5* (green), *S100β* (white) for astrocytes (**A-B**), and *NeuN* (cyan) for neurons (**C-D**), counterstained with the nuclei-dye Hoechst (blue), which provides the nucleus contour (white dashed line). Scale bar: 2 µm. (**B**, **D**) Bar chart of the mean *Fkbp5* mRNA puncta within the Hoechst contour with SEM. Each dot represents one contour, 20 cells per animal, 3 animals/condition. (**B**) Kruskal-Wallis test. ^ns^*p* > 0.05, not significant. (**D**) Kruskal-Wallis with selected Dunn’s multiple comparisons post hoc test, **p < 0.01, ***p < 0.001.

**Supplementary Figure 7.**
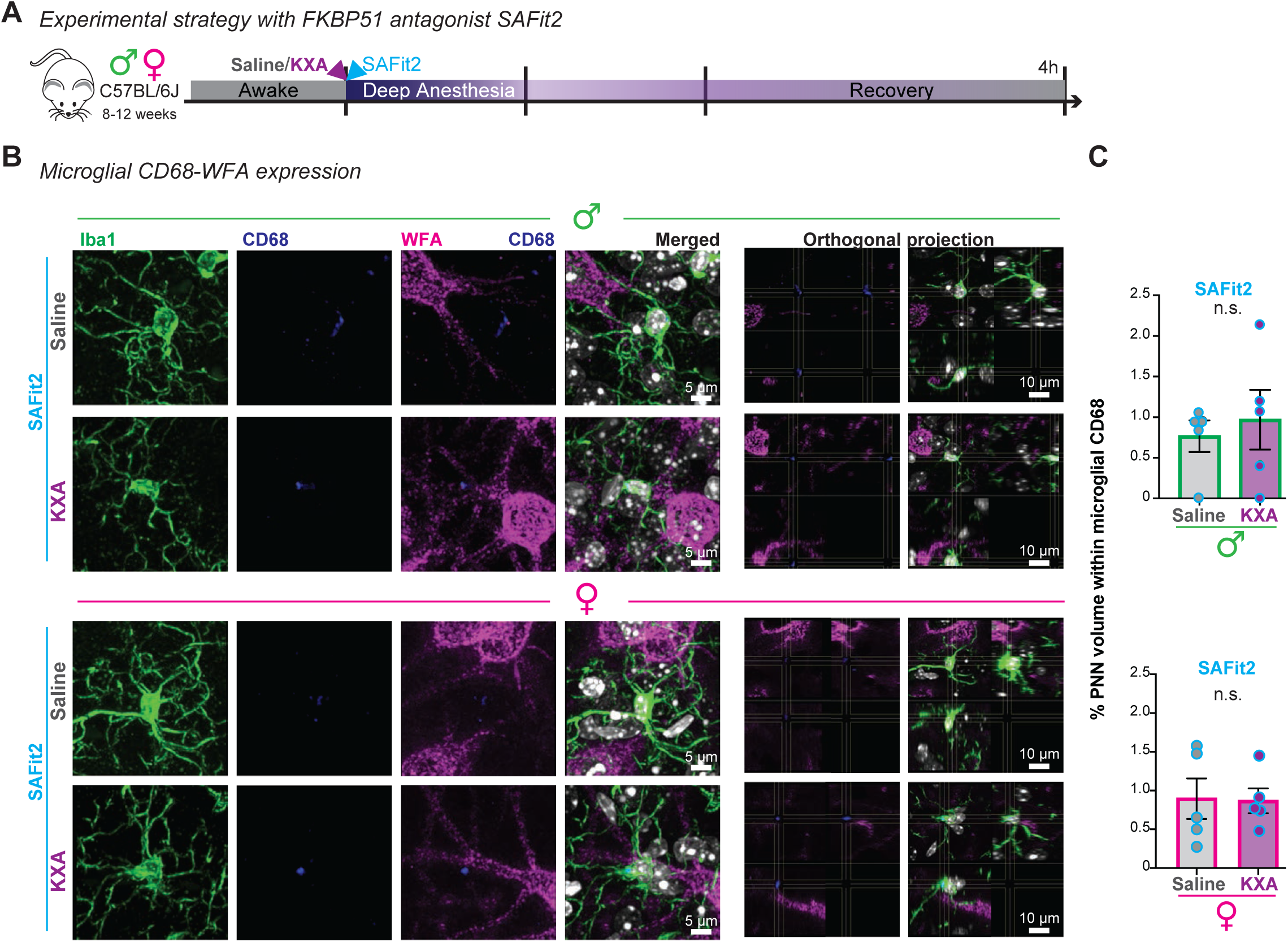
SAFit2 treatment inhibits female microglia from remodeling the perineuronal nets (PNN). (**A-C**) Antagonising FKBP51 with SAFit2 after injection of saline or KXA (ketamine-xylazine-acepromazine). (**B-C**) Comparison of perineuronal nets (PNN) staining inside microglia CD68. (**B**) Representative images of immunostained microglia with Iba1 (green), CD68 (blue), *Wisteria floribunda agglutinin* (WFA, magenta) for perineuronal nets (PNN), and counterstained with the nuclei-dye Hoechst (white) in the primary visual cortex (VISp), layer III-V of males and females, 4 hours after saline or KXA injection. Scale bar: 5 µm. Next to the merged image, orthogonal projections. Scale bar: 10 µm. (**C**) Bar charts of the mean percentage of PNN volume within microglial CD68 between males (green, top) and females (magenta, bottom) with SEM. Each dot, one animal. 5 animals/condition. Males (top bar chart), Unpaired t test with Welch’s correction. Mann-Whitney test and Unpaired t test with Welch’s correction. Females (bottom bar chart), Unpaired t test with Welch’s correction. Mann-Whitney test and Unpaired t test with Welch’s correction. ^ns^*p* > 0.05, not significant.

**Supplementary Figure 8.**
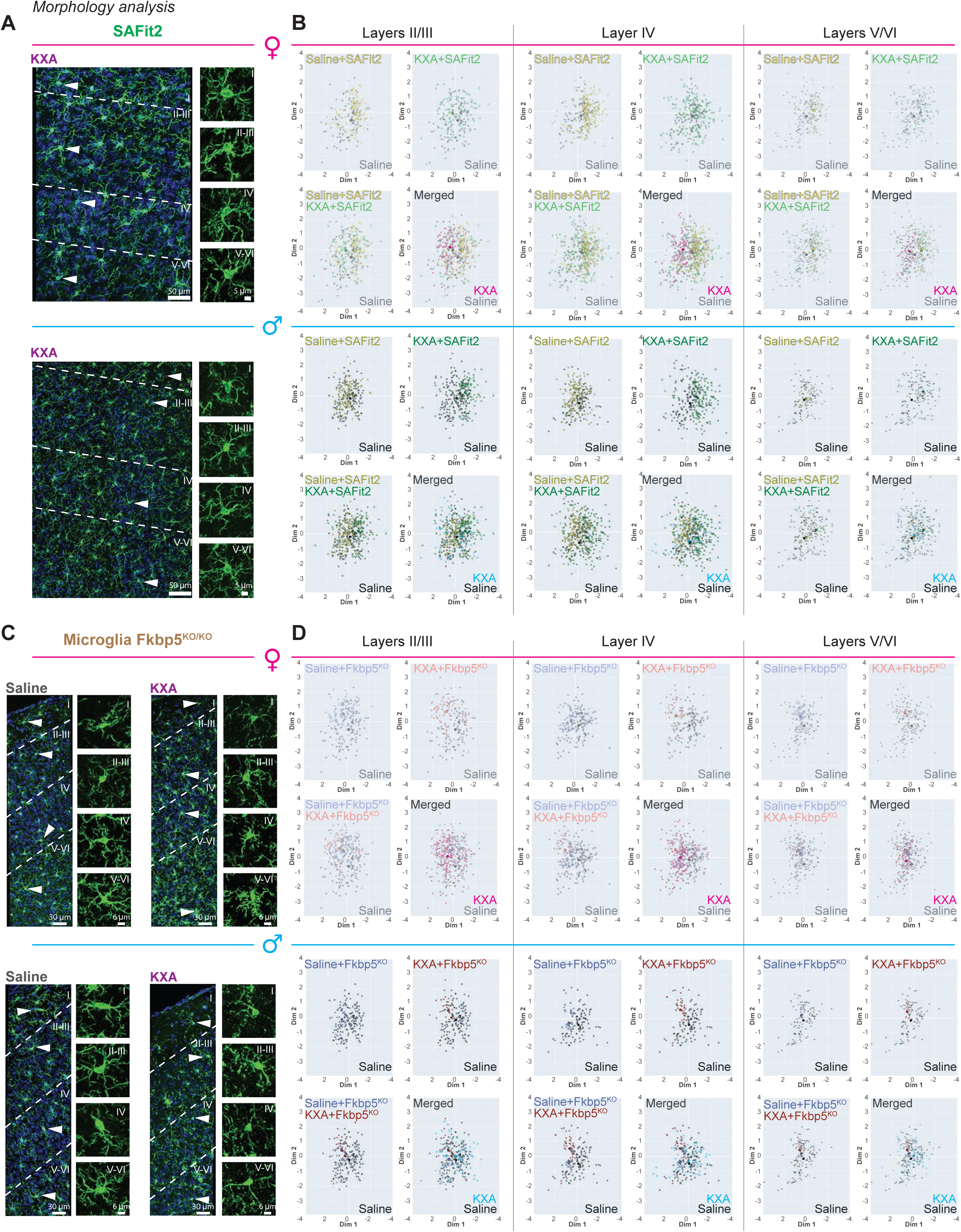
Morphological analysis of microglia across VISp layers after SAFit2 or microglia-specific *Fkbp5*^KO^. Morphological analysis of microglia after saline or KXA (ketamine-xylazine-acepromazine) with (**A-B**) antagonising FKBP51 with SAFit2 after injection in C57BL/6J or (**C-D**) in microglia-selective *Fkbp5*-knockout experiment using a tamoxifen-inducible Cx3cr1^CreERT2/-^ crossed with *Fkbp5*^KO/KO^ reporter mouse line (Microglia Fkbp5^KO/KO^). 3 consecutive tamoxifen injections, starting 8 days before performing the procedure. (**A**, **C**) Representative immunostainings for Iba1 (green) and CD68 (magenta) counterstained with the nuclei-dye Hoechst (blue) in the primary visual cortex (VISp) of females (magenta) and males (cyan). Left, overview image of the cortical layers. Scale bar: 50 µm (**A**), 30 µm (**C**). Arrow in each layer, microglia chosen for zoom-in. Scale bar: 5 µm. (**B**, **D**) Morphological analysis of SAFit2 (**B**) and microglia *Fkbp5*^KO/KO^ (**D**) microglia morphology within cortical layer II/III, IV, and V/VI in VISp after saline (grey, black) or KXA (magenta, cyan) using morphOMICs. Each microglia’s persistence image was embedded in a latent space using a Variational Autoencoder (VAE, see Methods). Larger dots, population mean for sex and condition.

**Supplementary Figure 9.**
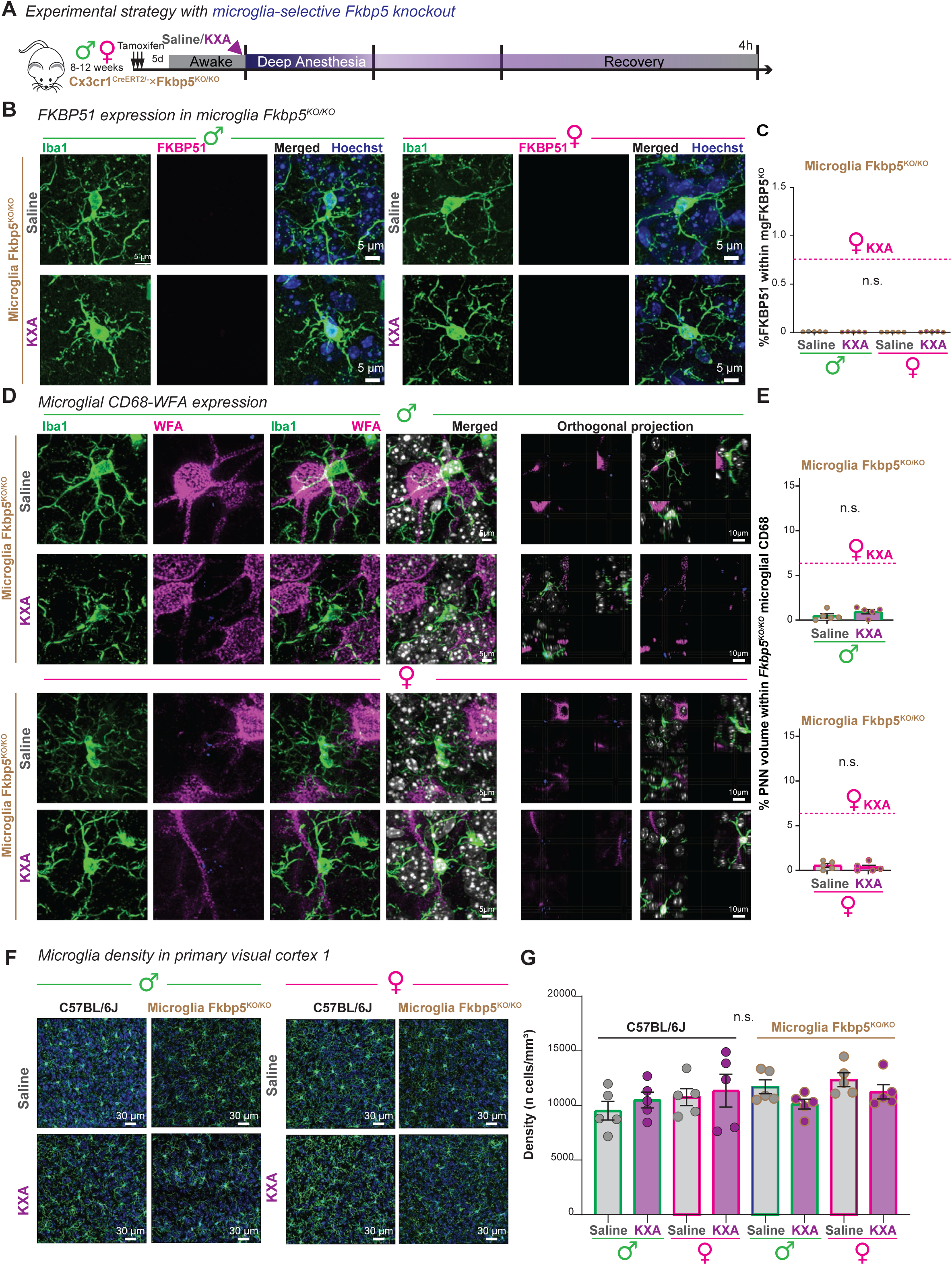
Microglia *Fkbp5*^KO^ prevents FKBP51 upregulation and PNN remodeling, without affecting microglia numbers. (**A-G**) Consequences of microglia-selective *Fkbp5*-knockout experiment in the primary visual cortex (VISp), layer III-V of males and females, 4 hours after saline or KXA injection using a tamoxifen-inducible Cx3cr1^CreERT2/-^ ×Fkbp5^KO/KO^ reporter mouse line (Microglia Fkbp5^KO/KO^). 3 consecutive tamoxifen injections, starting 8 days before performing the procedure (**A**). (**B-C**) Confirmation of FKBP51 knockdown. (**B**) Immunostaining for FKBP51 protein expression (magenta) in microglia (Iba1, green), counterstained with the nuclei-dye Hoechst (blue). Scale bar: 5 µm. (**C**) Bar chart of the mean percentage of FKBP51 volume within microglia with SEM. Each dot, one animal, 5 animals/condition. Magenta line, reference value of female KXA for FKBP51 (Figure 2G). Two-way ANOVA. ^ns^*p* > 0.05, not significant. (**D-E**) Comparison of perineuronal nets staining inside microglia CD68 between males (green) and females (magenta). (**D**) Representative images of immunostained microglia with Iba1 (green), CD68 (blue), *Wisteria floribunda agglutinin* (WFA, magenta) for perineuronal nets (PNN), and counterstained with the nuclei-dye Hoechst (white) in the primary visual cortex (VISp), layer III-V of males and females. Scale bar: 5 µm. Next to the merged image, orthogonal projections. Scale bar: 10 µm. (**E**) Bar charts of the mean percentage of PNN volume within microglial CD68. Each dot, one animal. 5 animals/condition. Magenta line, reference value of female KXA for PNN within microglia CD68 (**Figures S2D**, **S2F**). Two-way ANOVA. ^ns^*p* > 0.05, not significant. (**F-G**) Microglia density. (**F**) Immunostaining for Iba1 (green), counterstained with the nuclei-dye Hoechst (blue). Scale bar: 30 µm. (**G**) Bar chart of the mean microglia density in the primary visual cortex (VISp), layer II-V, across conditions. Each dot, one animal. 5 animals/condition. Kruskal-Wallis test. ^ns^*p* > 0.05, not significant.

**Supplementary Figure 10.**
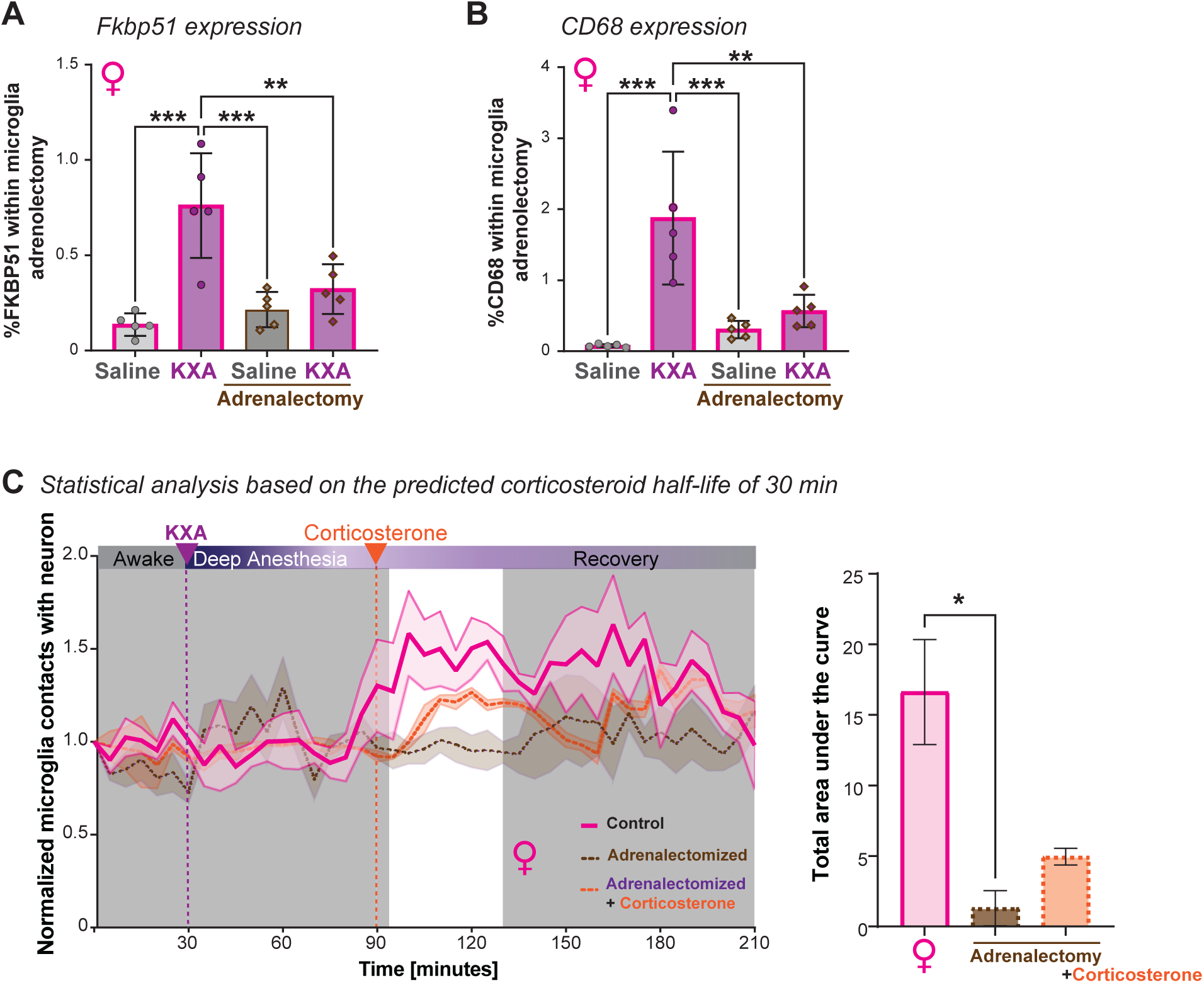
Comparison of FKBP51 and CD68 expression level in microglia in control and adrenalectomized females. (**A-B**) Bar chart of the mean percentage of FKBP51 (**A**) and CD68 (**B**) volume within adrenalectomized female microglia with SEM. Each dot, one animal, 5 animals/condition. Two-way ANOVA with selected Tukey’s multiple comparisons post hoc test, **p < 0.01, ***p < 0.001. (**C**) Normalized number of microglia and Thy1-EGFP neuronal process contacts over time in females (magenta, see Figure 1B), adrenalectomized females (purple, n=3), and adrenalectomized females injected with corticosterone 60 minutes after KXA injection (orange, n=3) represented as mean ± SEM confidence band. Dashed lines: KXA (green) and corticosterone (orange) injections. White band, area used for statistical analysis based on predicted corticosteroid half-life in rodents. Next, bar chart of the mean total area under the curves in (**C**) (magenta female from Figure 1C). Mean ± SEM of 3 - 5 animals per condition. Brown-Forsythe ANOVA test with Dunnett’s T3 multiple comparisons test, *p < 0.05.

**Supplementary Figure 11.**
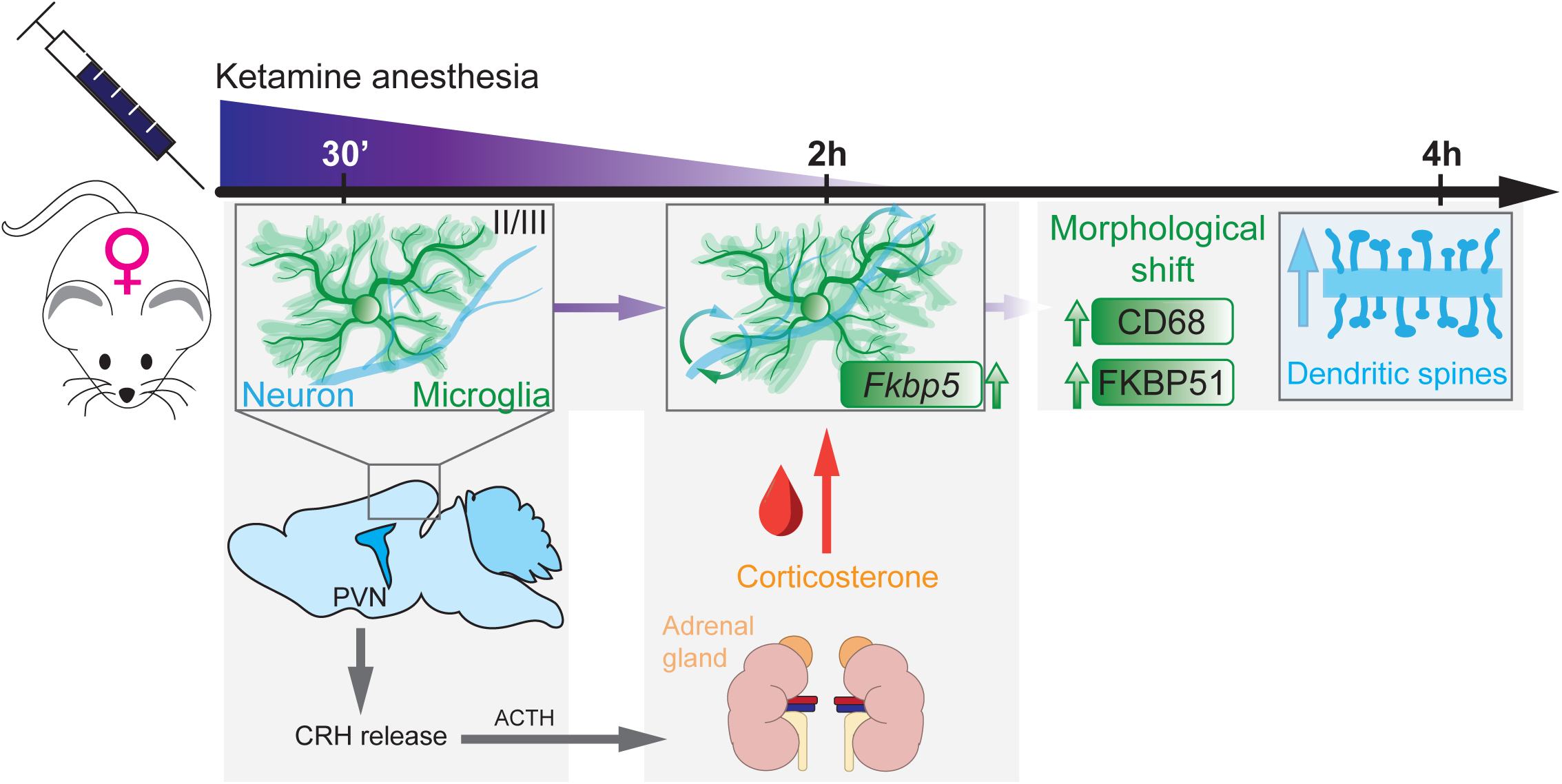
Summary figure. Schematic overview about the effects of ketamine anesthesia during the recovery on microglia (green) and neurons (blue). II/III, cortical layer II and III. ACTH, adrenocorticotropic hormone. CRH, corticosterone-releasing hormone. PVN, paraventricular nucleus.

## Notes

### Competing Interest Statement

The authors have declared no competing interest.

